# Adhesive and mechanical properties of the glue produced by 25 Drosophila species

**DOI:** 10.1101/2024.05.08.593221

**Authors:** Manon Monier, Jean-Noël Lorenzi, Sunitha Narasimha, Flora Borne, Vincent Contremoulins, Louis Mevel, Romane Petit, Youssef El Hachem, François Graner, Virginie Courtier-Orgogozo

**Affiliations:** Université Paris Cité, CNRS, Institut Jacques Monod, 75005 Paris, France; SMILE group, Center for Interdisciplinary Research in Biology (CIRB), Collège de France, 75006 Paris, France; Department of Biological Sciences, Columbia University, New York City, New York, USA; Inserm, B3OA, 75010 Paris, France; Université Paris Cité, CNRS, Matière et Systèmes Complexes, 75013 Paris, France

**Keywords:** bioadhesion, Drosophila, glue, attachment, adhesion assay, pupal shape

## Abstract

Drosophila glue, a bioadhesive produced by fly larvae to attach themselves to a substrate for several days, has recently gained attention for its peculiar adhesive and mechanical properties. Although Drosophila glue production was described more than 50 years ago, a general survey of the adhesive and mechanical properties of this proteinaceous gel across Drosophila species is lacking. To measure adhesion, we present here a protocol that is robust to variations in protocol parameters, pupal age and calculation methods. We find that the glue, which covers the entire pupal surface, increases the animal rigidity and plasticity when bound to a glass slide. Our survey of pupal adhesion in 25 Drosophilidae species reveals la wide range of phenotypes, from species that produce no or little glue and adhere little, to species that produce high amounts of glue and adhere strongly. One species, *D. hydei*, stands out from the rest and emerges as a promising model for the development of future bioadhesives, as it has the highest detachment force per glue area and produces relatively large amounts of glue relative to its size. We also observe that species that invest more in glue tend to live in more windy and less rainy climates, suggesting that differences in pupal adhesion properties across species are shaped by ecological factors. Our present survey provides a basis for future biomimetic studies based on Drosophila glue.

## Background

A fruitful approach to solve complex problems is to draw inspiration from nature, known as biomimetism. In particular, new bioadhesive materials are currently being developed for wet surfaces and medicine based on mussel glue (1). In comparison, natural glues that work in dry environments are less well known. The glue produced by Drosophila flies before metamorphosis stands out as a promising model for biomimetic dry adhesion (for review see (2)). In *Drosophila melanogaster* the glue is made of several long, glycosylated mucin proteins and short, non glycosylated proteins. It accumulates within the salivary glands during the last larval instar and is expelled by the animal just before entering metamorphosis. Once in contact with air, the glue dries out in a few minutes and allows the animal to remain attached to a substrate for several days, until the adult goes out of the pupal case. After glue expectoration, the cuticle hardens through a process named pupariation and the animal becomes a prepupa. Then, at 4-6 hours after puparium formation (APF), the epidermis progressively separates from the larval cuticle, starting anteriorly (3). At 11-13 h APF, the head everts, the oral armature of the larva is detached and the animal technically becomes a pupa (4, 5). For simplicity, and because the exact developmental stage is not always known in our experiments, we name the animal, from pupariation to adult eclosion, a “pupa”. Given the large range of genetic tools available and its rearing ease, *D. melanogaster* appears as a great model to study glue characteristics, e.g. by introducing mutations in glue genes or by expressing exogenous glue genes.

More than 1500 species of Drosophila genus (6) have been collected all over the world, including a few cosmopolitan ones and mostly endemic species. Drosophila species live in diverse habitats across all climatic conditions: deserts, forests, mountains, caves, buildings (7). Since Drosophila pupae are 1-2 mm long, brown and immobile, they are difficult to spot in the wild and relatively little is known about their ecology. For a few species such as *D. suzukii*, pupae do not appear to be fixed to a substrate and lie several centimetres deep into the soil (8). For other species, pupae have been found attached to a wide variety of substrates: the dry parts of various rotten fruits and cacti, grape stalks, wood, glass bottles (2, 9). We can thus hypothesise that the different Drosophila glues have been optimised and adapted to a wide range of conditions, and that they display varying adhesive and mechanical performances.

Glue production and composition has been examined in a dozen of Drosophila species (2), but no general survey of glue adhesive and mechanical properties across Drosophila has been undertaken. We previously developed a method to measure Drosophila glue adhesion, published in 2020 (10). As it is not yet possible to extract the glue and carry out direct experiments on it, we are obliged to measure adhesion of the pupa. Briefly, late wandering larvae are placed on glass slides within a plastic box and by the next day all animals are attached to the glass slide (or to the plastic of the box). The glass slide is then fixed with holders onto a platform. A force sensor covered with double-sided sticky tape goes down until it touches the animal, then the sensor goes up at constant speed and at some point the pupa is detached. Force is measured across the entire experiment. The adhesion force is the maximum force at which detachment occurs. For *D. melanogaster* adhesion force ranges between 151 mN and 269 mN for 1.1 mm^2^, giving an adhesion strength of 137–244 kPa (10), which corresponds to some of the strongest bioadhesives (11). Compared to daily life adhesives, Drosophila glue is thus positioned between repositionable adhesives (∼10 kPa) and neopren/cyanoacrylate glues (∼10^3^/10^4^ kPa) glues, like strong adhesive tapes (∼100 kPa).

Several substrates of various roughnesses, hydrophilic and charge properties were tested and all displayed similar adhesion forces (around 200 mN) except teflon (40 mN). Intraspecific variation was also assessed using 12 *D. melanogaster* strains from diverse geographical regions (12). The surface of glue in contact with the substrate was not significantly different between strains but median adhesion force varied between 120 and 375 mN. Regarding interspecific variation, adhesion force was reported for three species besides *D. melanogaster, D. simulans* (234 mN), *D. suzukii* (78 mN) and *D. hydei* (482 mN), all on glass slides; all three exhibited the same adhesion strength (13).

Here, we use our previously published experimental set up (10) to measure pupal adhesion in various species and we broaden our past analyses by assessing other mechanical properties besides adhesion force, in particular elasticity and plasticity. Our aim is to examine pupal adhesion in multiple Drosophilidae species that can be bred in the laboratory and to identify species with the best potential for developing future adhesives.

## Materials and Methods

### Fly culture and stocks

Flies were cultured in plastic vials on standard medium (4 l: 83.5 g yeast, 335.0 g cornmeal, 40.0 g agar, 233.5 g saccharose, 67.0 ml Moldex, 6.0 ml propionic acid). For *D. suzukii, D. prostipennis, D. kurseongensis and D. rhopaloa* this medium was supplemented with 200 g of D-glucose anhydrous (VWR Chemicals, reference: 24379.294). Stocks, their origin and their temperature of culture are given in Table S1.

### Adhesion assays

The pull-off force necessary to detach the pupa from the glass slide was measured using a universal test machine (LS1S/H/230 V Lloyd Instruments) with a 5N force sensor (YLC-0005-A1 Lloyd Instruments), as described previously (12). Third instar wandering larvae were washed in PBS to remove traces of food and microorganisms from their surface. They were then dried by placing them briefly on tissue paper and transferred on glass slides (Menzel Superfrost microscope glass slide, ThermoScientific™ n° AGAB000080) with soft forceps within a closed box that was maintained on humid wet cotton at 25°C (for all species). Between 15 and 21 hours after transfer, pupae naturally attached to the glass slides with their ventral part adhering to the glass slide, and not in contact with other pupae, were used for adhesion measures.

In the standard protocol, we used Tesa, extra strong, n° 05681-00018 tape as double-sided adhesive tape. The force sensor covered with the double-sided adhesive tape was moved down with a constant speed of 0.016 mm/s until it pressed the pupa with a force of 0.07 N. It was then stilled at a force of 0.03 N for 10 s and finally moved up with a constant speed of 0.2 mm/s until the pupa was detached. The force, time and sensor position were recorded using NEXYGENPlus software (Lloyd Instruments). We also used alternative protocols, which corresponded to the standard protocol with one or two parameters modified (Table S2).

For *D. simulans, D. yakuba* and *D. mauritiana*, adhesion tests experiments were retrieved from previous work (12), which used the standard protocol.

### Classification of adhesion results

After adhesion assay, three scenarios are observed for the pupa (Table S3-4): (a) The pupa detaches from the glass slide, and remains on the piece of double sided tape on the sensor; (b) The pupa is broken in two parts. In this case, the head remains attached onto the glass slide while the remaining body is found on the double sided tape on the sensor. The break usually occurs along the operculum, which is detached and stays onto the sensor while the part of the head and the anterior pupal case in contact with the slide remain onto the slide; (c) The pupa is not detached after adhesion assay, the full body remains on the glass slide and is not damaged.

### Analysis of adhesion tests data

Scripts written on R version 4.1.2 (2021-11-01) were used to analyse adhesion test data and prepare figures. Some figures were then modified using Inkscape 1.2.1 (2022-07-14 version).

We note *F*_*i*_ the force (in N) and *P*_*i*_ the sensor position/extension (in mm) at landmark i. *F*_*i*_ is positive when the sensor presses onto the pupa and is negative when the sensor pulls on the pupa. *P*_*i*_ is always positive and increases when the sensor goes down towards the pupa (as in the beginning of the experiment).

Seven landmarks are defined and positioned on the force and distance recordings (Fig. 1). There are several ways to automatically label these landmarks and here is what we chose for the general case. For each given experiment, we first estimated the noise before contact (before B), noted σ, as the standard deviation of the force calculated over the first measurements (representing 4% of the measurements recorded during the experiment). The noise after detachment (after G), noted α, was defined as the amplitude range of the last recorded values of the force (representing 10% of all values). Landmark A corresponds to the beginning of the experiment. In region AB the sensor is going down. Landmark B is the moment at which the sensor comes into contact with the pupa. In this general case, in region BC, when the sensor presses onto the pupa, the force goes above 3σ for more than 3 seconds. Landmark B is defined as the moment when the force exceeds σ, just before going above 3σ for more than 3 seconds. (Checking that the force goes above 3σ for more than 3 seconds allows to eliminate spurious force peaks due to disruptive events such as a shock to the table). In region BC the sensor is pressing onto the pupa until reaching a given force (0.07 N for standard protocol, or else 0.25 N, see Table S2) at the maximal position. Landmark C is when this lowest position is reached. Then the protocol applies a constant, smaller constant force (0.03 N in the standard protocol, or else 0.21 N) for a given time (10 s in the standard protocol, else 0 or 5 min), from landmark D (first time the force is constant) to landmark E (end of the compression phase). In region EFGH the sensor is going up. The force decreases, becomes null at landmark F, goes below -α and then reaches the maximal negative force (the detachment force). When the detachment force is lower than −3α, we consider that there is a detachment peak and G is defined when the force increases and goes above -α. Landmark H is the end of the experiment. In region GH the sensor is going up, carrying away the free pupa, and is no longer pulled down.

**Figure 1.**
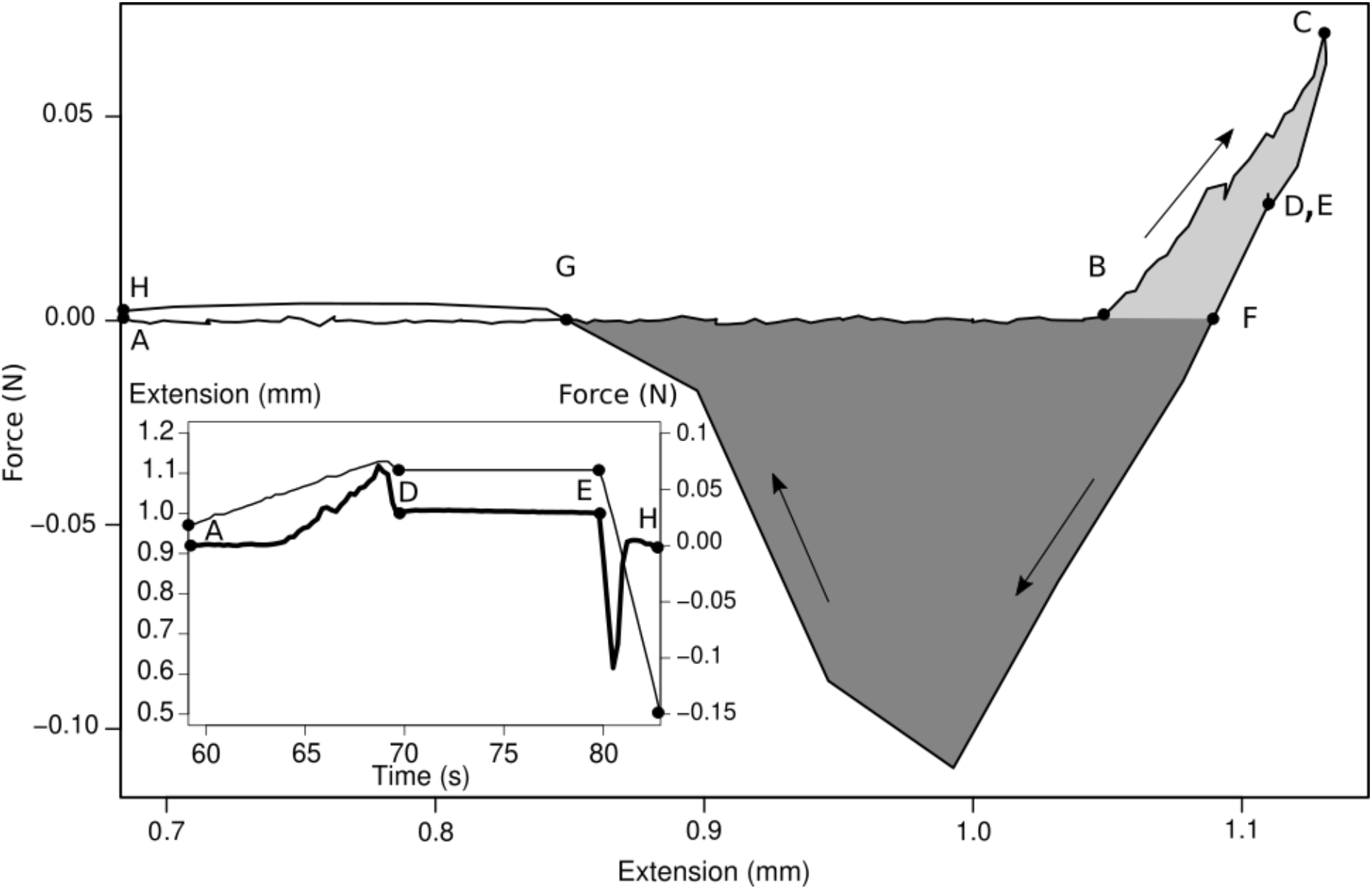
One example of force and distance recordings obtained with our pupa adhesion assay (here for *D. melanogaster* Canton S with the standard protocol). Main diagram: force versus extension; arrows indicate the direction of time flow. Inset: extension (thin line) and force (thick line) versus time. Seven landmarks are positioned along the curves. A: beginning of the experiment, B: first contact between sensor and pupa, C: maximal force applied to the pupa, D: a constant force starts to be applied to the pupa, E: the sensor starts going up, F: the force is null, G: the pupa is detached from its substrate, H: end of the experiment. See text for details. On the main diagram, landmarks D and E coincide. In the inset, some landmarks have been omitted for clarity. The one-way detachment energy (DE1) corresponds to the dark grey area and the two-way detachment energy (DE2) to the sum of the light and dark grey areas.

Beside the general situation explained above, we distinguished several other cases. In situations where σ was incorrectly estimated (e.g. the sensor touches the pupa before recording, shock to the machine, etc.), we were not able to define landmark B as above and considered that landmarks A and B colocalized. In the situation where protocol ‘5 min’ was used to press the pupa, our machine created higher noise in the position values, so we defined landmark C at the maximal force of the experiment. In the situation where protocol ‘0 s’ was used (to press the pupa), the time between landmark B and landmark C was less than one second and landmark C was defined at 0.5 seconds after landmark B. In situations where the force after landmark E did not decrease below −3α (most often because the pupa was poorly attached), landmark G was positioned when the force value first reached -α after landmark F.

From these force and position recordings one can estimate five mechanical quantities relating to the pupa and its glue: the detachment force, the one-way and two-way detachment energies (DE1 and DE2, respectively), the deformation (Def) and the rigidity, as explained below. The detachment force is the extremum force measured when the pupa detaches. The one-way detachment energy (DE1) is the area of the negative part of the decompression curve: 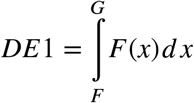. The two-way detachment energy (DE2) is the area between the compression curve and the decompression curve: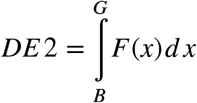.

The deformation (Def) of the animal and its glue is the difference of position between the first contact with the pupa (landmark B) and the last contact with the pupa (landmark G): Def=*P*_*B*_*-P*_*G*_. It is equal to zero when the deformation is fully reversible. It is positive when the pupa stretches more than in its resting state, and negative when the pupa remains compressed when stretched relative to its resting state. The rigidity (or stiffness, noted *k*) represents the resistance of an object to deformation and is obtained here by dividing the force applied to the pupa by its deformation upon compression: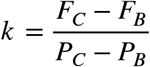.

During the adhesion assays, air humidity, air temperature and atmospheric pressure were recorded so that we could test their potential effects on adhesion measures. Using all data on 25 species, including pupae that were either detached, not detached or broken during adhesion assays, we first tested the effect of stocks using a one-way ANOVA with the R aov function (force ∼ Stock). We found that most of the variation in adhesion force is explained by the variable “Stock” (p<2e-16, 44% of the variance explained). Using a four-way ANOVA, we found that the variables “Humidity”, “Temperature” and “Atmospheric pressure” are not independent of the variable “Stock” (File S7). This relates to the fact that our adhesion tests were performed on different days for the various fly stocks and species. In order to test the effect of the variables “Humidity”, “Temperature” and “Atmospheric pressure”, we then tested statistical differences in adhesion forces for D. melanogaster data only (343 adhesion measures) using the R aov function (force ∼ Humidity*Temperature*Pressure) and detected no significant effect of air humidity, air temperature and atmospheric pressure (File S7).

### Phylogenetic trees

Phylogenetic trees representing the evolution of adhesion properties through evolution were computed using Mesquite software (version 3.8 http://www.mesquiteproject.org/). Phylogenetic tree topology is from (14). Adhesion force median values for each species are given to the software, which infers the ancestral state for each branch, based on a parsimony reconstruction method. Adhesion force is illustrated by a colour gradient attributed to branches.

### Measurements based on pupae pictures

To assess pupal size and shape and the surface of glue in contact with the glass slide, three images of the pupa on the slide were taken before adhesion tests with a Keyence microscope VHX-2000 Z20 ×20 on dorsal, ventral and side views. Note that adhesion tests were first done without taking pictures and that for some species or some animals, pupa pictures are not available. From the dorsal view images two values were automatically extracted: the pupal area and the pupal maximal length. First, an ImageJ (version 1.53c) macro converted the image files into a .tif format in order to be processed by Ilastik (version 1.3.2 https://www.ilastik.org/) software. Second, these converted images were used for machine learning pixel classification in Ilastik. We used the pixel classification function to allocate pixels to two object classes: the pupa vs the rest of the image. The features used to distinguish these two classes are all the features available in Ilastik regarding colour and intensity, edge characteristics and texture parameters. Ilastik uses the machine learning algorithm Random Forest classifier, which is interactively trained from user annotations. The experimenter manually selects pixels on the image with two different colours: one to label the pupa and another to label the rest of the picture. Based on these manual selections, the Random Forest algorithm assigns a probability for each pixel of the image to belong to the pupa or the rest of the picture. To facilitate the training process, pictures were manually separated into two groups: light pupae and dark pupae. Training was separately performed on 6% pictures of each group of pictures (i.e. 70 light and 30 dark pupae) randomly chosen between different fly species and different protocol conditions.

Ilastik automatically classified pixels for the other images. According to the probability obtained for each pixel, binary masks were generated by Ilastik segmenting the pupa and the rest of the image. In cases where two pupae were adjacent and merged in the binary mask, we manually separated them by drawing a thin line between them with the pencil tool in ImageJ. Lastly, the pupal area and the pupal length (defined as corresponding to the maximum Feret diameter) were calculated with a custom-made ImageJ macro. Based on these measures, we evaluated pupal shape as 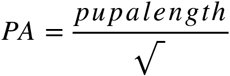.

Ventral view pictures were used to measure glue area, which is defined as the surface of glue in contact with both the glass slide and the animal. Pictures were first anonymized and then a single person (RP) manually outlined and measured the glue print areas using ImageJ. Based on these pictures, we evaluated glue investment as the dimensionless number 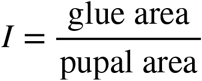 and adhesion strength as 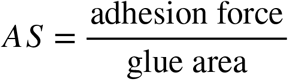. Side view pictures were used to measure pupal height manually in ImageJ, i.e. the maximal distance between the ventral side and the dorsal side, perpendicular to the glass slide.

### Collection and analysis of ecological data

We analysed the meteorological data of the collection sites for the studied Drosophila species. For seven species, we used the latitude and longitude indicated by colleagues who had collected or received the flies (Table S1). For the other species, for which only the country or region of collection is known, we used the latitude and longitude of the most central location of the country or region, using the ‘Coordonnées GPS’ website (https://www.coordonnees-gps.fr/). We provided a list of the latitude and longitude for each location to the Open Meteo website (https://open-meteo.com/en/docs/historical-weather-api/#location_mode=csv_coordinates) and retrieved daily weather data over the last 10 years (from 02/25/2014 to 02/25/2024). Open Meteo website used estimated latitude and longitude, within a 25 km resolution. We obtained the elevation, maximum, minimum, mean temperature and apparent temperature, the daylight and sunshine duration, rain, snowfall, precipitation (total of rain, snowfall and ice rain), wind speed, wind gusts, the shortwave radiation sum and the reference evapotranspiration (File S7). We retrieved the number of cells per salivary gland from (15). We extracted the mean, minimal and maximal values over the last 10 years for each meteorological parameter. Scripts written on R version 4.1.2 (2021-11-01) were used to analyse meteorological test data and prepare figures.

### Humidity preference in laboratory conditions

Three pieces of Whatman paper (Fisher Scientific, 11728742) cut into 5.5 cm diameter circles were placed in a 12 cm x 12 cm square Petri dish (Greiner, n° 688102) at equal distance from each other, without contact and attached with double-sided tape (Tesa extra strong, n° 05681-00018) (Fig. S23). The first paper circle was left dry, the second wetted with 200 µL of fresh tap water and the third with 400 µL. About 40-60 third instar wandering larvae were placed with entomological forceps in the middle of the Petri dish. Petri dishes were closed and placed in a large plastic box (36 cm x 24 cm x 14 cm) whose bottom was covered with wet paper. An empty Petri dish was placed at the bottom of the large box. The large box containing Petri dishes was then placed in an incubator overnight, at 25°C and 80% of humidity. Between 15 and 21 hours after transfer, pupae were found to be naturally attached within the Petri dishes. We counted the number of pupae attached to the three Whatman papers with different levels of humidity and to the plastic of the Petri dish. For each species, we observed 3-4 Petri dishes over four different days and measured the fraction *f*_*0*_ of total pupae attached on dry paper circles, *f*_*200*_ and *f*_*400*_ on wet ones and *f*_*plastic*_ on the plastic Petri dish. The fractions *f*_*0*_, *f*_*200*_, *f*_*400*_ and *f*_*plastic*_ are numbers between 0 and 1 and their sum is one. The humidity preference index (a value between −1 and 1) was calculated as the difference (*f*_*400*_-*f*_*0*_). In total, we tested the humidity preference of 21 species. For *D. eugracilis, D. pachea, D. kurseongensis* and *D. prostipennis* we did not obtain enough larvae.

### Statistical tests

Statistical tests were performed in R version 4.1.2 (2021-11-01).

## Results

### Our measure of pupal adhesion is not sensitive to protocol parameters

To test whether the assay we developed previously to detach pupae and assess adhesion force (10) can affect the measure of pupal adhesion, we carried out variations of our standard protocol by changing only one or two parameters at a time, using Canton S *D. melanogaster* pupae (Table S2). We found that the detachment force was not affected by the maximal force at which the pupa was pressed (0.07 N or 0.25 N), the time during which the pupa was pressed (0 s; 10 s; 5 min), the speed at which the sensor was moved during the compression or decompression phases (speed x3 or speed/3) or the type of double-sided tape used (Fig. S1). This indicates that our measure of adhesion force is robust and does not change with these experimental parameters.

In our standard protocol, the adhesion test is performed 15-21 hours after wandering L3 larvae are deposited on glass slides. To test whether pupal age can affect adhesion measures, we performed adhesion tests 3 days later, i.e. 87-93 hours after wandering L3 larvae were deposited on glass slides. We found that the adhesion force was not affected by pupal age (Fig. S1).

The detachment force, which represents the maximal force measured at the time of pupa pulling off, was the unique measure of adhesion that we assessed in our previous studies (Fig. 1) (10, 12, 13). Other measurements of the adhesion are possible. We examined two other ones, the energy necessary to detach the pupa and the energy necessary to detach the pupa and to press it beforehand, as per our protocol. Both measurements were calculated as the areas delimited by the force-position curves, the first one only during the decompression phase (named one-way detachment energy, DE1), and the second one during both the compression and decompression phases (named two-way detachment energy, DE2) (Fig. 1). Across all protocols, we found that the detachment force, the one- and the two-way detachment energies were correlated (Fig. S2-3) and were not affected by modifications of the protocol parameters, including changes in pupal age (Fig. S2). Overall, our measures of pupal adhesion appear to be robust to variations in protocol parameters, pupal age and methods used to estimate the adhesion.

### The glue increases the animal rigidity and plasticity

Since our adhesion assays comprise both a compression and a decompression phase, the force-distance-time curves allowed us to assess the animal rigidity and plasticity. First we verified that the standard protocol is adapted for rigidity calculation: the rigidity is the slope of the force-position curves and these curves are effectively linear when the maximal applied force is 0.07 N (Fig. S4). We also noted that when we use the 0.25 N protocol, the rigidity increases beyond 0.08 N due to the non-linearity of the force-position curves (Fig. S4) and therefore we did not measure rigidity with this 0.25 N protocol. Overall, we found that our measures of rigidity are robust to variations in protocol parameters: the time during which the pupa was pressed (0 s; 10 s; 5 min), the speed at which the sensor was moved during compression or decompression phases (speed x3 or speed/3) and the tape used on the sensor (Tesa or Gergonne) (Fig. S5). The rigidity of the animal (corresponding to the pupa and the glue onto it) is higher in adhesion tests performed three days later (Fig. S5A), and this is consistent with the hardening of the pupal cuticle over time (16).

Comparing the force-time curves between the loading and unloading phases (from landmark B to C and from D to F) shows that attached pupae are plastic: their deformation is significantly irreversible. This irreversible contribution is unchanged when the speed of compression or decompression is increased or decreased by 3 and hence is not of viscous origin (Fig. S5B). Our measures of plasticity are also robust to most changes in protocol parameters: the tape used on the sensor (Tesa or Gergonne) and the time during which the pupa is pressed (0 s; 10 s). However, when the animal is pressed for 5 min, it gets more deformed than with the standard protocol. Interestingly, plasticity is unchanged for three-days older pupae (Fig. S5B).

In addition to its contact with the solid substrate, the glue covers the ventral part of the pupal surface, as shown by scanning electron microscopy (10), so we wondered whether the glue could affect the animal rigidity and plasticity. We compared attached pupae with pupae that were gently detached with a brush from the glass slide and with detached pupae that were sticked back on a piece of Tesa tape. We found that detached pupae placed on a glass slide or a piece of Tesa tape are less rigid than pupae attached with their own glue (Fig. S5A). Thus the glue, when in contact with a solid substrate, appears to increase the effective animal rigidity. Furthermore, detached pupae are elastic (their deformation is fully reversible), whereas naturally attached pupae and pupae glued back on Tesa tape are plastic. After compression a pupa can thus go back to its initial shape within seconds if detached and its residual deformation appears to be due to its adhesion to the glass slide (via tape or via its own glue).

### Pupae from several species do not detach after adhesion assay

To evaluate the diversity of glue adhesion forces in flies, we performed our standard adhesion assay on 24 other Drosophilidae species (Table S1), spanning approximately 60 million years of evolution (17). For most species, almost all the processed pupae fully detached and did not break, as for *D. melanogaster* (Table S3, Fig. S6A). However, for six species (*D. virilis, D. pachea, D. nannoptera, D. immigrans, D. hydei* and *Zaprionus lachaisei*), more than 20% of the processed pupae did not detach from the glass slide or broke during the stretch phase (Fig. S6A). We then used a stronger tape (the strongest we could find) and increased the maximal force applied to the pupa (0.25 N) for these six species, and for another species, *D. littoralis*. This new protocol (‘1 strong tape; 0.25 N’) allowed us to detach all pupae from *Z. lachaisei* and a higher proportion of pupae for the other species (Fig. S6B-C, Table S4). We note that even with this strongest tape, a few pupae of these other species (48% at most) broke or did not detach from the glass slide. The median detachment force measured with the ‘1 strong tape; 0.25 N’ protocol was comparable to the median detachment force measured with the standard protocol for all species except for *D. hydei* where it was stronger (Fig. S7). In the next analyses, we thus pooled data from both adhesion protocols for all species except *D. hydei*, for which we used only the ‘1 strong tape; 0.25 N’ protocol, which gave stronger adhesion values. The adhesion force measured for undetached or broken pupae was slightly higher than for detached pupae (Fig. S8), suggesting that the breakage and absence of detachment occur with higher adhesion forces. Given that for undetached or broken pupae the measured detachment force may be lower than the actual adhesion, we examine below only detached pupae. Overall, our adhesion assays are adapted to all the fly species we tested except *D. hydei, D. virilis, D. littoralis, D. nannoptera* and *D. pachea*, for which they provide a lower estimate of their adhesion force. For these species, even with the most adhesive tape commercially available, 15-25% of the pupae do not detach (Table S4).

### *D. hydei* and *D. pachea* display the highest adhesion strength

To test whether the adhesion force is influenced by pupal size, pupal shape and the surface of glue in contact with the glass slide and the animal (named “glue area”), we examined pictures of the animal from dorsal, ventral and side views that were taken before the adhesion assays for 17 selected species (Fig. S9-10). Pupal length and height increase with pupal area, with *D. virilis* and *D. littoralis* being significantly more elongated than other studied species (Fig. S11). For some pupae, we could not detect any glue area on the ventral pictures (Figs. S9, S12). The proportion of pupae with no visible glue area is about 80-85% for *D. tropicalis* and *D. rhopaloa*, 20-45% for *D. malerkotliana, D. prostipennis* and *Z. indianus*, and 5-10% for *D. funebris, D. nannoptera* and *Z. lachaisei* (Fig. S12). Median glue area tends to increase with pupal size (Fig. S13), *D. hydei, D. virilis* and *Z. lachaisei* being among the largest pupae and producing the most glue (Fig. 2).

**Figure 2.**
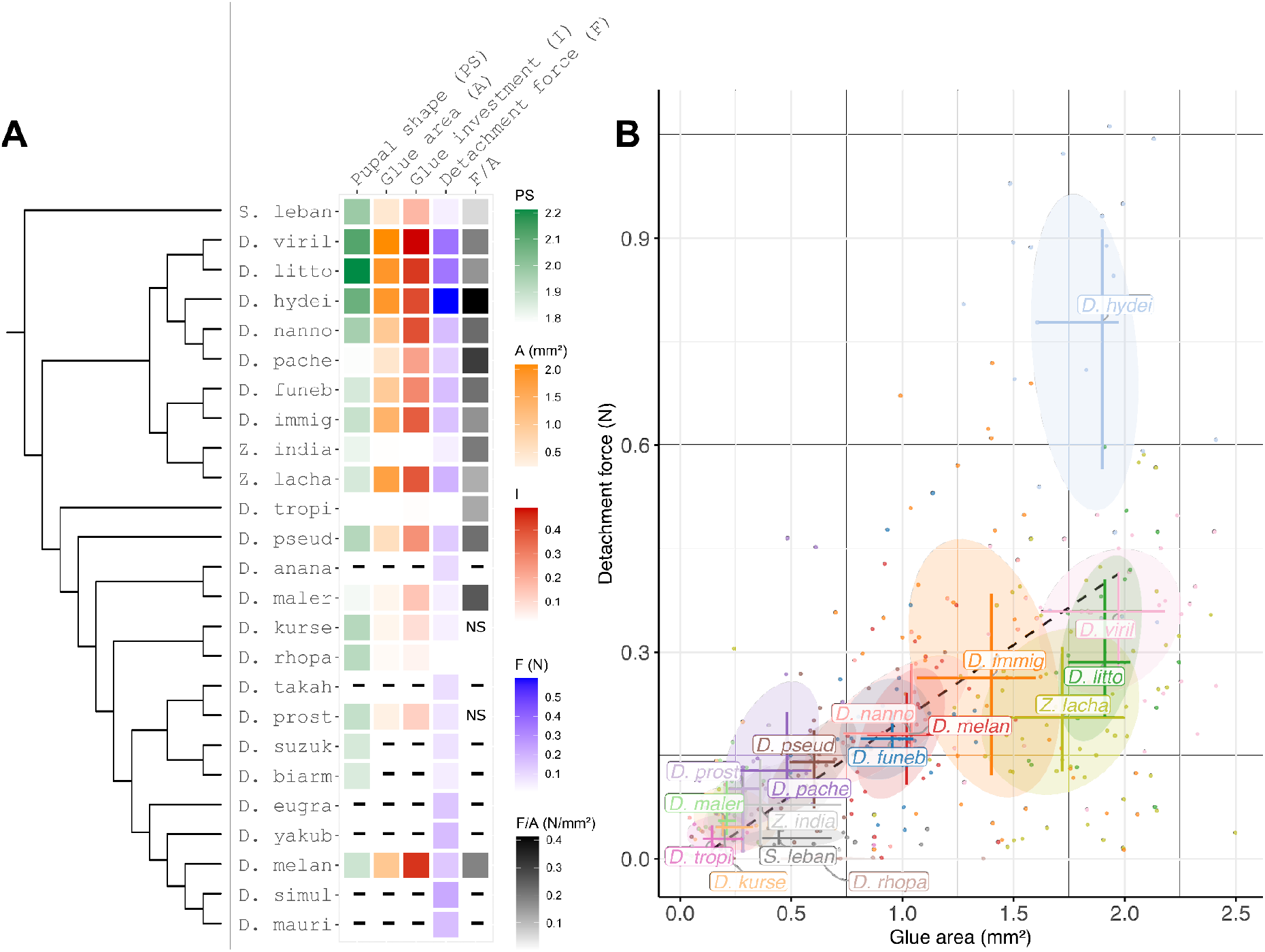
Geometrical and mechanical measurements of glue in various Drosophila species. (A) MMedian pupal shape, glue area (mm^2^), glue investment, detachment force (N) and adhesion strength (N/mm^2^)). (B) Distribution of the various species across the range of detachment forces and glue areas. Each dot corresponds to a single pupa detached during the adhesion assay. The centre of the crosses corresponds to the median. Horizontal and vertical bars represent the first and third quartiles. The dotted line is the regression line between the median values for each species (r^2^=0.61, p<0.0002). Medians in A are represented with a colour code, light and dark colours respectively representing low and high values (pupal shape in green, glue area in purple, detachment force in blue and adhesion strength in grey). Hyphens represent unavailable data and ‘NS’ data not shown because less than 10 measurements were obtained. Species names are abbreviated using their first five letters. Phylogenetic tree is adapted from (14) and tree branch lengths do not represent real distances. PS: pupal shape, A: Glue area, I: Glue investment, F: Detachment force, F/A: adhesion strength (Detachment force/Glue area).

For all 25 tested species, the one- and two-way detachment energies (Fig. S14) correlates with the detachment force (Fig. S15), so we decided to use the detachment force as a simple and relevant measure of adhesion for the remaining part of our study. As expected, the higher the glue area, the higher the adhesion force (Fig. S16), but no clear correlation is found between detachment force and pupal size (Fig. S17). Interestingly, the adhesion force correlates with the surface of glue in contact with the pupa relative to the pupal area, named ‘glue investment’ (Fig. S18) and this correlation, r^2^=0.75, is almost as high as the correlation of pupal and glue areas, r^2^=0.85 (Fig. S13). Strongly attached species are generally the ones producing a large surface of glue relative to their body size. Furthermore, the adhesion force per unit of glue area (adhesion strength) is comparable between all 17 measured species except *D. hydei and D. pachea*, which have the highest median adhesion strength (Fig. 2 and Figs. S19-S21).

Taken together, our results suggest that adhesion strength is comparable between most species, except *D. hydei* and *D. pachea* whose glues are more adhesive and that the interspecific variation in pupal adhesion is mostly due to differences in the amount of glue produced.

### In more windy and less rainy climates flies tend to invest more in glue

To understand how differences in pupal adhesion properties across species may be shaped by ecological factors, we retrieved meteorological parameters from the sites where the various species were collected and evaluated their correlation with pupal adhesion properties. We found that species investing the most in their glue are the ones found in drier and more windy environments (Fig. 3). On the opposite, species with a lower glue investment are found in more humid and less windy environments. For other ecological parameters, no clear correlation with adhesion properties was found (Table S5, Fig. S22). Meteorological data obtained at collection sites may not always correspond to the local microclimate experienced by small animals such as flies. To further examine the relationship between glue investment and atmospheric humidity, we compared humidity preference for pupariation sites between species. We gave larvae a choice for pupariation between three papers of distinct moistures (Fig. S23). Among 21 tested species, we observed that *D. rhopaloa, D. virilis* and *Z. lachaisei* larvae show a strong preference for the highest humidity whereas *Scaptodrosophila lebanonensis* larvae choose the driest surface (File S7). Other species did not show such clear preferences. We did not observe any correlation between adhesion properties and humidity preference for pupariation sites between species (Table S5). Overall, the highest correlation was between glue investment and precipitation (Table S5, Fig. 3A).

**Figure 3.**
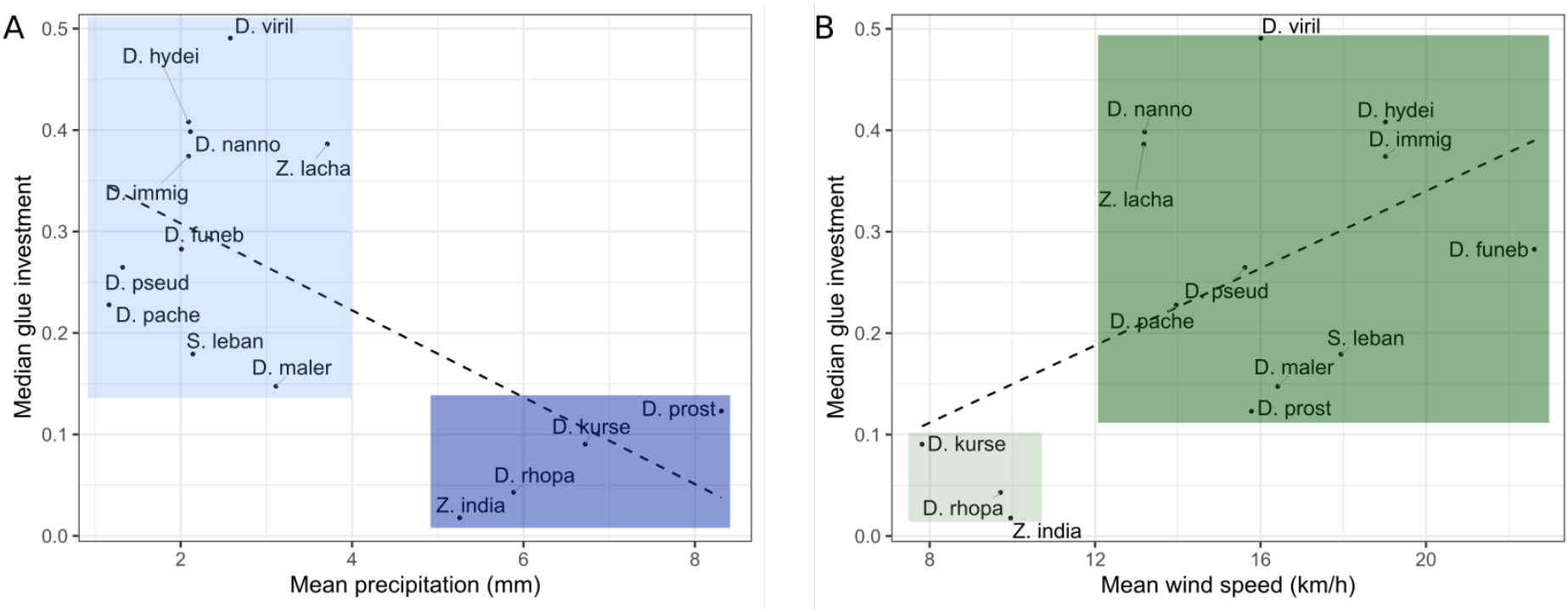
Glue investment versus precipitation and wind speed. Glue investment is estimated as glue area divided by pupal area. (A) Median glue investment versus mean precipitation (mm). Species collected in a rainier environment and with a lower glue investment are highlighted in dark blue and species from a drier environment investing more in glue are in light blue. (B) Median glue investment versus wind speed (km/h). Species from a more windy environment investing more in glue are highlighted in dark green and species collected in a less windy environment with a lower glue investment are in light green. Species names are abbreviated using their first five letters. Dashed lines are linear regressions: A: r^2^=0.4, p=0.016; B: r^2^=0.27, p=0.059.

## Discussion

In 2020 we published the first report – as far as we know – of an adhesion assay to measure the force necessary to detach a fly pupa from a substrate (10). Then we used our protocol to assess pupal adhesion on glass slides for 12 strains of *D. melanogaster* (12) and for 5 other Drosophila species (12, 13). Here, we show that our assay is robust and not affected by changes in protocol parameters. Whereas our protocol can detach all the *D. melanogaster* pupae, during this study we came across Drosophila species for which a significant proportion of the pupae did not detach with our assay, indicating that our tape was not adhesive enough. Then, using the most adhesive tape we could find, we managed to obtain a higher proportion of detached pupae but still, for *D. pachea* and *D. hydei* about 25% of the tested pupae did not detach (Table S4). While we were polishing up this manuscript, Beňo et al. published a paper (19) where they presented an adhesion assay similar to ours, except that 7.5-mg-metal rings are mounted on the dorsal side of the animal using Loctite SuperBond instant glue and that the adhesion test is performed two hours later after the glue is dry. They examined adhesion of one *D. melanogaster* line (Oregon-R) on glass slides and found adhesion forces and strengths comparable to ours (19). We previously tested such an application of Loctite Super Glue-3 onto the pupa to measure its adhesion but our trials were inconclusive: we couldn’t prevent the liquid glue from sliding along the pupal walls and reaching the glass slide, and the glue appeared to induce a reaction onto the cuticle, hardening it. Our present method, using double-sided sticky tape, can be more high-throughput, produces less waste and preserves the animal rigidity, allowing us to measure not only adhesion but also rigidity and plasticity.

Although we have sought to limit measurement noise in our experiments by working with standardised glass slides, uncrowded fly cultures and choosing only animals attached on their ventral side and not in contact with others, the adhesion forces we measured were relatively variable between individuals of the same species (for example, from 0 to 428 mN for D. melanogaster). Such a wide range of adhesion force values was also noted in previous reports by ours (10) and others (19). This variation can be due to individual variability in diverse factors such as pupal size, pupal shape, pupal position relative to the substrate, the amount of glue, the shape and surface of the glue contact area relative to the shape and size of the animal, and the possible presence of traces deposited by larvae on the glass slide before pupal attachment. In future studies it would be interesting to try to set up a novel adhesion assay using an entirely different method, for example by collecting the glue while it is expelled by the animal. We note that a centrifugal test such as in (15) would not be feasible as more than 20,000 g would be required to detach a 1-mg pupa.

In addition to the maximal force at detachment, we examined two other measures of adhesion, one-way and two-way detachment energies (DE1 and DE2). DE2 is the energy necessary to press the pupa and then detach it, as per our protocol, while DE1 corresponds to the energy necessary to detach the pupa. For pupae that are at least as adhesive as *D. melanogaster* we observed that the energy to press the pupa (before pulling) is negligible and that DE1 and DE2 are highly correlated and proportional for different protocols (Fig. S3A) and species (Fig. S14). DE1 and DE2 also correlate with the detachment force (Fig. S3B-C).

As a byproduct of our method, we measured the rigidity of *D. melanogaster* pupa. In line with the hardening of the cuticle over time (3), we observed a significantly higher rigidity three days later (Fig. S5A). Young modulus for *D. melanogaster* pupa is about 1 MPa (approximately 1 N/mm for a 1 mm thick pupa), which corresponds to that of polystyrene foam or rubber, and is thus a thousand times more than that of single typical epithelial cells (20). Interestingly, pupae attached with their own glue are more rigid than detached pupae (Fig. S5A). The glue may act as a splint when animals are attached to a rigid substrate such as a glass slide. Indeed, the glue not only lies at the contact surface between the animal and the substrate, but also covers the ventral surface of the pupa (10) and extends onto the lateral sides of the fly via bifurcated glue channels, named bidentia (19). Previously, we noticed that ants take a longer time (several dozens of minutes) to pierce attached pupae than detached pupae with their mandibles (13). This piercing is the first step for ants eating Drosophila pupae. The glue thus appears to protect flies from ant predation via two mechanisms, first by increasing pupal rigidity and making it difficult for the ant to grab or pierce the pupa, and second by fixing it strongly to a substrate so that ants cannot detach it and bring it to the nest.

Our examination of pupal adhesion in 25 Drosophilidae species reveals a continuous distribution of phenotypes, from non adhesive species to highly adhesive species. At the lowest are species that produce no or little glue and adhere little: D. tropicalis, D. rhopaloa, D. malerkotliana, D. prostipennis, D. kurseongensis and Z. indianus. For these species, our measures of glue adhesion strength, obtained by dividing the detachment force by small values close to detection limit, are not reliable. At the highest are species that have large pupae, produce the highest quantities of glue relative to their size and have the most adhesive pupae: D. littoralis, D. virilis and D. hydei. Overall, the variation in adhesion between species seems to be mostly related to differences in glue investment and glue area (Fig. S13). One species, D. hydei, stands out as a particularly promising model for the development of future bioadhesives, as it has the highest detachment force per glue area (Fig. S2).l Note that we examined only one strain per species, hence the extent of intraspecific variation in adhesion, outside *D. melanogaster* (12), remains unknown. Furthermore, we only assessed adhesion on glass slides. In the future, we plan to test other substrates and various temperature/humidity conditions.

Our results reveal that *D. virilis* and *D. hydei* display the strongest adhesion force and the highest surface of glue, suggesting their salivary glands produce larger amounts of glue than other species. Oddly, their salivary glands contain fewer cells than in *D. melanogaster* (82 and 110 cells per gland, respectively, versus 134 for *D. melanogaster*) (15). A recent study (15) reported the number of cells per salivary gland for ten of the species analysed here. Overall, we found no correlation between the number of these secretory cells and the glue area or glue investment (Table S5). This indicates that most of the variation we observed between species in the amount of glue produced is not explained by changes in the number of secretory cells. Other factors that may influence the quantity of glue are external environmental conditions (substrate, temperature, etc.), larval nutritional conditions, the degree of secretory cell polyploidy, secretory cell size, secretion rate, duration of the secretory phase,, as well as the quality of glue expulsion behaviour (21).

To calculate glue adhesion strength, we divided the detachment force by the glue area that is in contact with both the pupal case and the glass slide (Fig. S10). However, other choices are possible. The glue forms a meniscus and extends both onto the pupal case and the glass slide (10). Furthermore, the glue extends anteriorly in alluvial-like layered terraces onto the glass slide (19) and is often partly mixed with a granular liquid probably expelled by the anus at the moment of glue expectoration (10). For substrates such as Teflon where all the glue is taken off with the puparium, Beňo et al. proposed to divide by the entire area of the substrate covered by the glue (19). For glass slides, only part of the glue in contact with both the pupal case and the glass slide usually goes away together with the pupa upon detachment (10). So our choice may be the most appropriate for our case. We note that we did not consider the thickness of the glue at the interface between the pupal case and the substrate, which may vary between species.

Given the rapid evolution of glue genes in Drosophila compared to other genes (9, 12, 22) and the large range of ecological niches occupied by the 25 Drosophila species examined in this study, we were expecting to find more conspicuous differences in adhesion between species than what we observed here. This suggests that the glue may have additional functions during the life of the animal. The extremely rapid rate of evolution of the glue genes, in terms of gene copy number and number of repeats and repeated motifs, suggests their involvement into an evolutionary “arms race” (23). Following glue release, the salivary glands produce an exuvial fluid that lies between the animal and its pupal case (24). This fluid contains antimicrobial and antibacterial factors and appears to protect the metamorphosing animal from pathogens (24). In fact, several observations suggest that the glue may also have immune functions, besides its adhesive and stiffening properties uncovered in the present study (2). First, the glue lies not only at the interface between the animal and the substrate but also appears to cover the ventral surface of the animal, as observed by scanning electron microscopy (10). Second, Drosophila glue contains several mucin-like proteins, and mucins are known to protect from microbial invasion (25). Third, yeast-like organisms and coliform bacteria firmly entrapped within the glue have been observed by scanning electron microscopy (26). Fourth, the glue protein Eig71Ee/gp150 is produced by both salivary glands and hemocytes, and participates in the entrapment of bacteria in the hemolymph (27). So far, little is known in *D. melanogaster* about antibacterial peptides and immunity at the prepupa/pupal stage compared to the larval and adult stages (28). Further work is needed to decipher the exact role of Drosophila glue in immunity.

During metamorphosis the animal is immobile and vulnerable, so that its ability to stick to a substrate can be of high importance for survival. The variation in adhesion that we detected between species, although smaller than expected, may reflect adaptation to distinct environments and pupariation sites. To investigate this hypothesis, we searched for correlations between adhesion parameters and meteorological factors at the places where the different species were collected. We found no correlation with glue area or glue adhesion. Interestingly, we observed that in more windy and less rainy climates flies tend to produce more glue relative to their size. The wind may, directly or indirectly, detach the animal and bring it to an unfavourable place for metamorphosis, with a different humidity, more exposure to predators, etc. In more windy climates larvae may thus have evolved to produce more glue relative to their size. In addition, as Drosophila researchers are well aware, a drop of water can facilitate the detachment of the puparium or pupa from the substrate (see also (19)). More rainy climates may select for other means of protection than glue adhesion. We note however that three species with distinct glue investment, *D. hydei, D. simulans* and *D. suzukii* were collected at the same spot (13). This confirms that the correlation we observed with meteorological factors is weak and that other factors probably influence glue-related traits.

Diptera species provide good models to study adhesion, as these organisms can be easily raised in laboratory conditions, are widely studied in genetics, and secrete a strong, biodegradable and biocompatible bioadhesive that can last for several days at least. Studying Diptera glue can have future applications in industry and may lead to the development of new adhesives that are safe for human health and the environment. Our analysis reveals that *D. virilis* and *D. hydei* are the most promising species to investigate further as they display the strongest adhesion force and they produce relatively large amounts of glue. In *D. virilis* three glue gene orthologs have been annotated (2) and in *D. hydei* we found five orthologs. Our study paves the way for a genetic analysis of their glue and should lead to the identification of genes and other characteristics that allow the formation of a highly adhesive biomaterial. The powerful genetic tools available in Drosophila and the diversity of ecological niches to which this fly glue adapted during evolution promise major advances in our understanding of complex biomaterials.

## Abbreviations

APF: after puparium formation
DE1: one-way detachment energy
DE2: two-way detachment energy
Def: deformation

## Acknowledgements

We thank P. Andolfatto, A. Farlow, P. Gibert, N. Gompel, M. Lang, S. Prigent and B. Prud’homme for flies. We also thank I. Nuez and S. Boutaleb for helping with fly work, S. Prigent for his help in examining the ecological characteristics of the various studied species, L. Le Boudec for Figure 2B, C. Gay for discussions about adhesives and reviewers for comments. We acknowledge the ImagoSeine core facility of the Institut Jacques Monod, member of the France BioImaging infrastructure (ANR-10-INBS-04) and GIS-IBiSA.

## Funding

This work was supported by CNRS as part of the MITI interdisciplinary action “Défi Adaptation du vivant à son environnement” and from the European Research Council under the European Community’s Seventh Framework Program (FP7/2007-2013 Grant Agreement no. 337579) to VCO. MM was supported by a PhD fellowship from “Ministère de l’Education Nationale, de la Recherche et de la Technologie” (MENRT) obtained from the BioSPC doctoral school. JNL was supported by the Labex “Who AM I?”, ANR-11-LABX-0071 and the Université Paris Cité, Idex ANR-18-IDEX-0001, funded by the French Government through its “Investments for the Future” program.

### Use of Artificial Intelligence (AI) and AI-assisted technologies

Random Forest algorithm for image segmentation.

## Data availability

Data is provided within the manuscript, in supplementary figures and in supplementary information files available at DRYAD: https://doi.org/10.5061/dryad.79cnp5j3j.

The code is also available at GitHub: https://github.com/manonmonier2/Adhesion/releases/tag/v1.0.0.

Final DRYAD link (will be active upon manuscript acceptance): https://doi.org/10.5061/dryad.79cnp5j3j

## Author contributions

V.C.-O., M.M. and F.G. conceived the project. V.C.-O. and F.G. supervised the project. M.M., S.N., F.B., L.M., R.P. performed the experiments. M.M., J.-N.L., F.G. and V.C.-O. examined the data. M.M., J.-N.L., V.C.-O. and Y.E.H. analyzed the data with scripts and prepared figures. V.C.-O. and M.M. wrote the original draft with input from F.G. All authors reviewed the manuscript. V.C.-O. acquired funding.

