## Supplemental Figures and Tables for "Adhesive and mechanical properties of the glue produced by 25 Drosophila species"

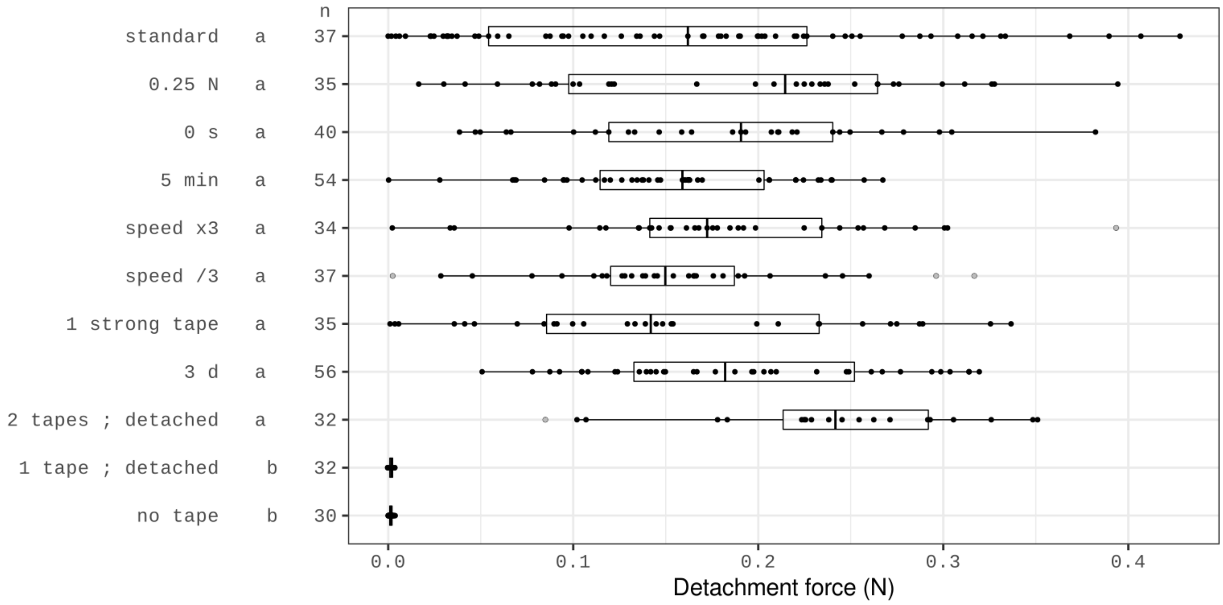


Figure S1. Detachment force (in N) for control adhesion assays with *D. melanogaster.* Each dot corresponds to a single pupa. Ends of the boxes define the first and third quartiles. The black vertical line represents the median. The horizontal line on the right of the box extends to the largest value not further than 1.5 × IQR (IQR, interquartile range is the distance between the first and the third quartiles) from the upper hinge of the box. The horizontal line on the left of the box extends to the smallest value not further than 1.5 × IQR of the hinge. Points outside of the horizontal lines are considered as outliers and plotted in grey. Vertical axis, left column: protocol name. The various protocols are described in Table S2. Middle column: an ANOVA followed by all pairwise comparisons after Tukey correction ($p < 0.05$) was performed on protocol groups. Protocols that are not significantly different from each other share a letter (e.g. a, b). Right column: *n* indicates the total number of pupae tested for each condition.

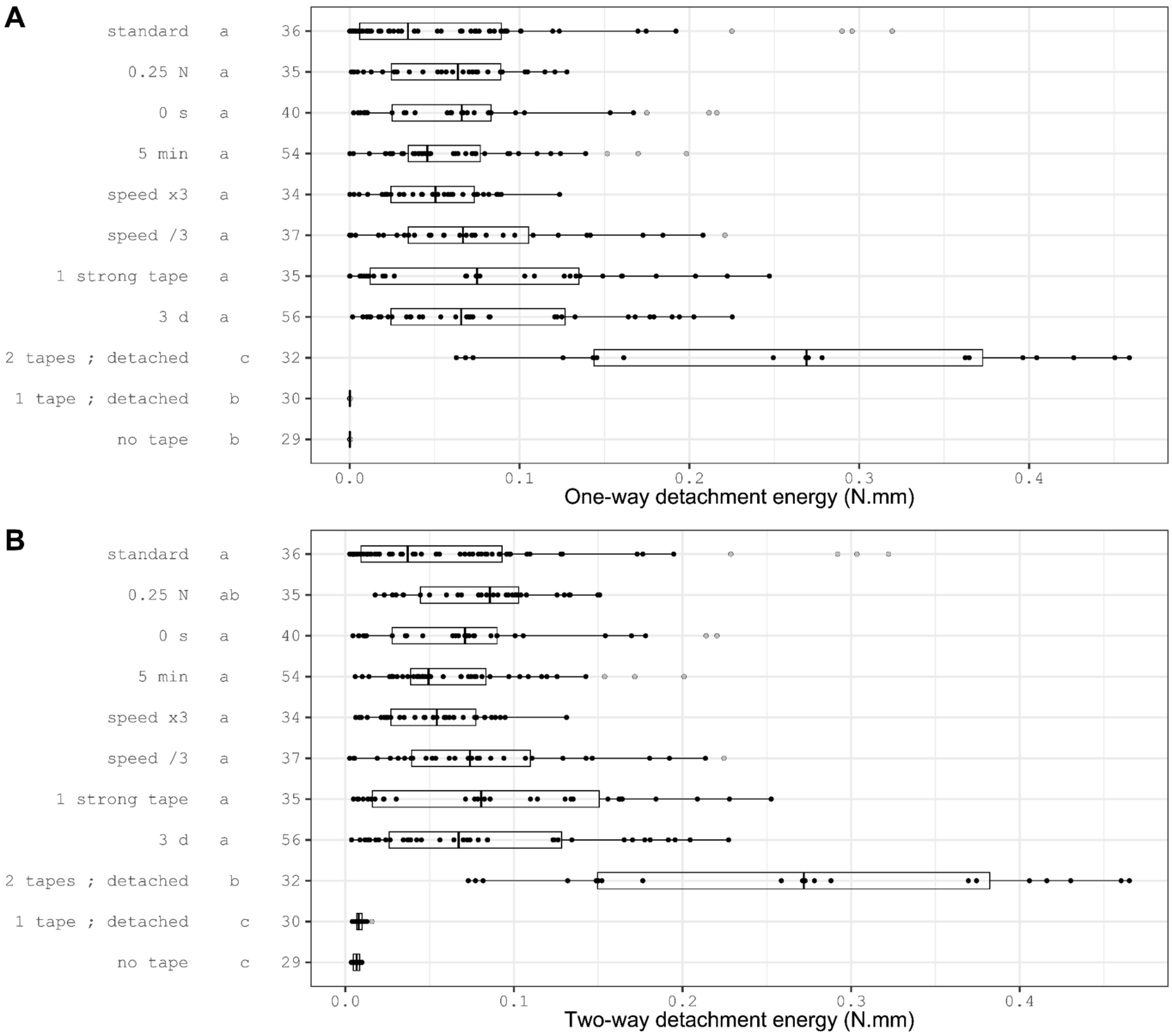


Figure S2. One-way (A) and two-way (B) detachment energy for control adhesion assays with *D. melanogaster.* Same legend as Fig. S1. The various protocols are described in Table S2.

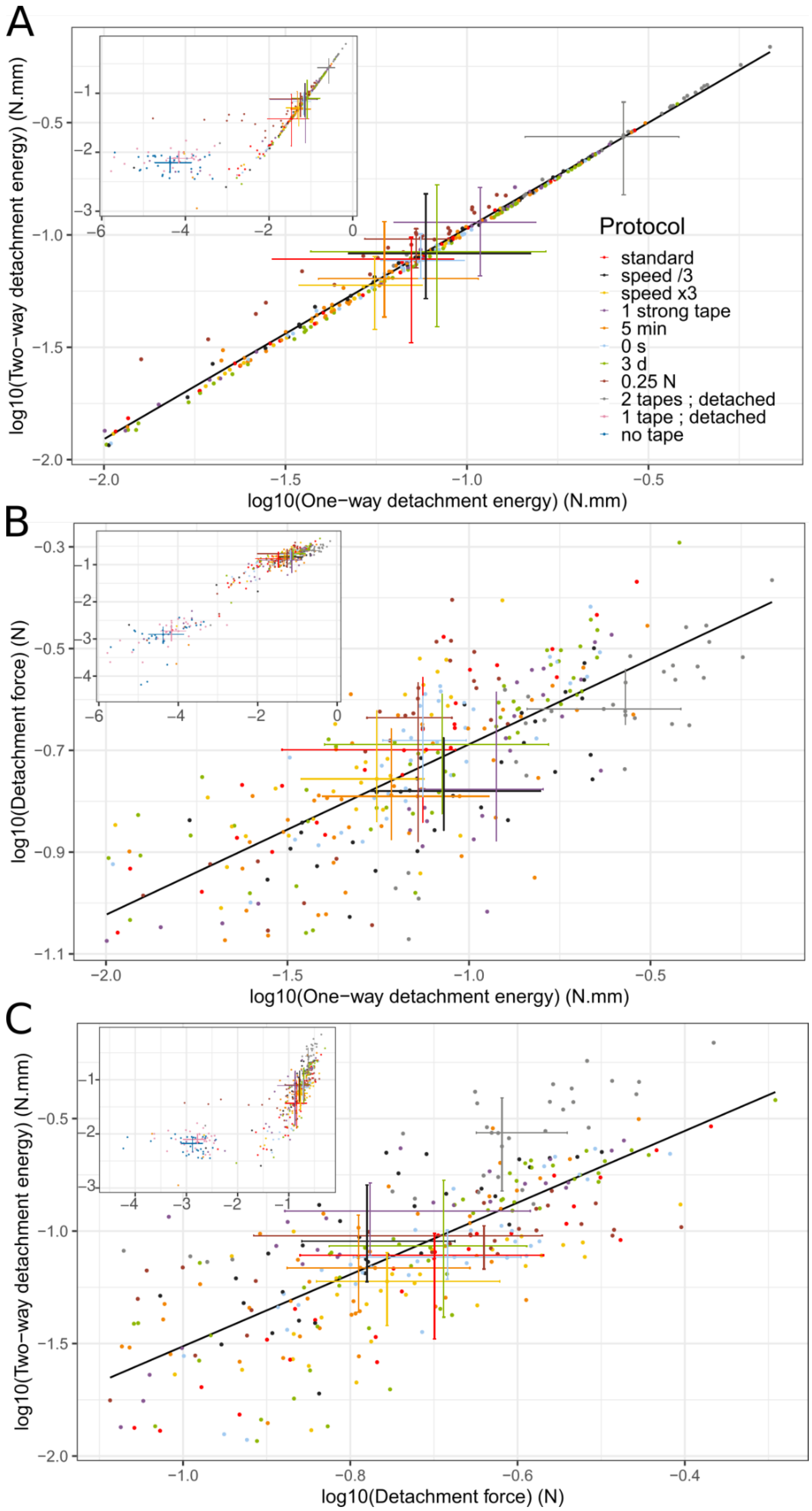


Figure S3. Detachment force and energies for control experiments conducted on Canton S *D. melanogaster*. (A) Two-way detachment energy versus one-way detachment energy. (B) One-way detachment energy versus detachment force. (C) Two-way detachment energy versus detachment force. Note the logarithmic scales. Main graphs are zooms of the inserted graph. Regions under 0.03 N are not displayed on zoomed graphs. Protocols used for control experiments are distinguished with colours. Each dot corresponds to a single pupa, either detached, broken or undetached after adhesion assay. The regression line using single pupa data points above 0.03 N is drawn in black. Center of crosses correspond to the median for each protocol. Horizontal and vertical bars represent first and third quartiles. Regressions: A: $r^2 = 0.99$, $p < 2.2e\text{-}16$ ; B: $r^2 = 0.56$, $p < 2.2e\text{-}16$ ; C: $r^2 = 0.56$, $p < 2.2e\text{-}16$.

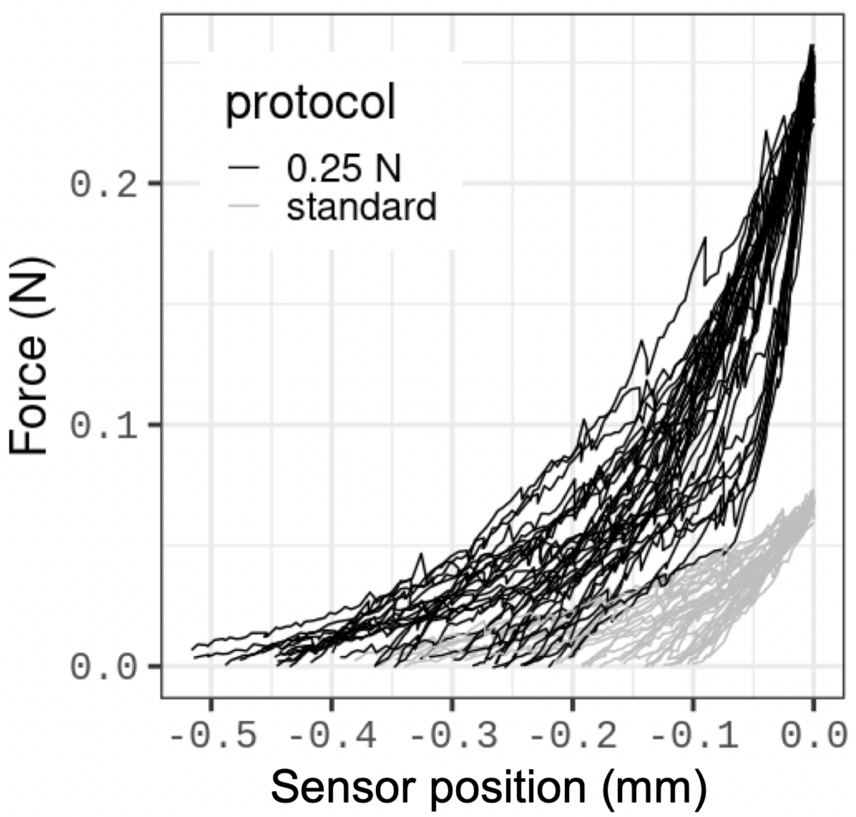


Figure S4. Pupal rigidity assessment for *D. melanogaster*. The force (in N) measured during sensor movement versus sensor position (in mm) between landmarks B and C (Fig. 1) is shown for the standard protocol (grey) and for the 0.25 N protocol (black) for 72 pupae in total. Sensor position is the position of the sensor, also named extension in the manuscript. For clarity, curves are aligned at the position of the maximal applied force, corresponding to landmark C. Rigidity corresponds to the slope of these curves. Curves corresponding to standard protocols are overall linear whereas the ones corresponding to 0.25 N tend to display two slopes, with a breakpoint between 0.07 N and 0.1 N.

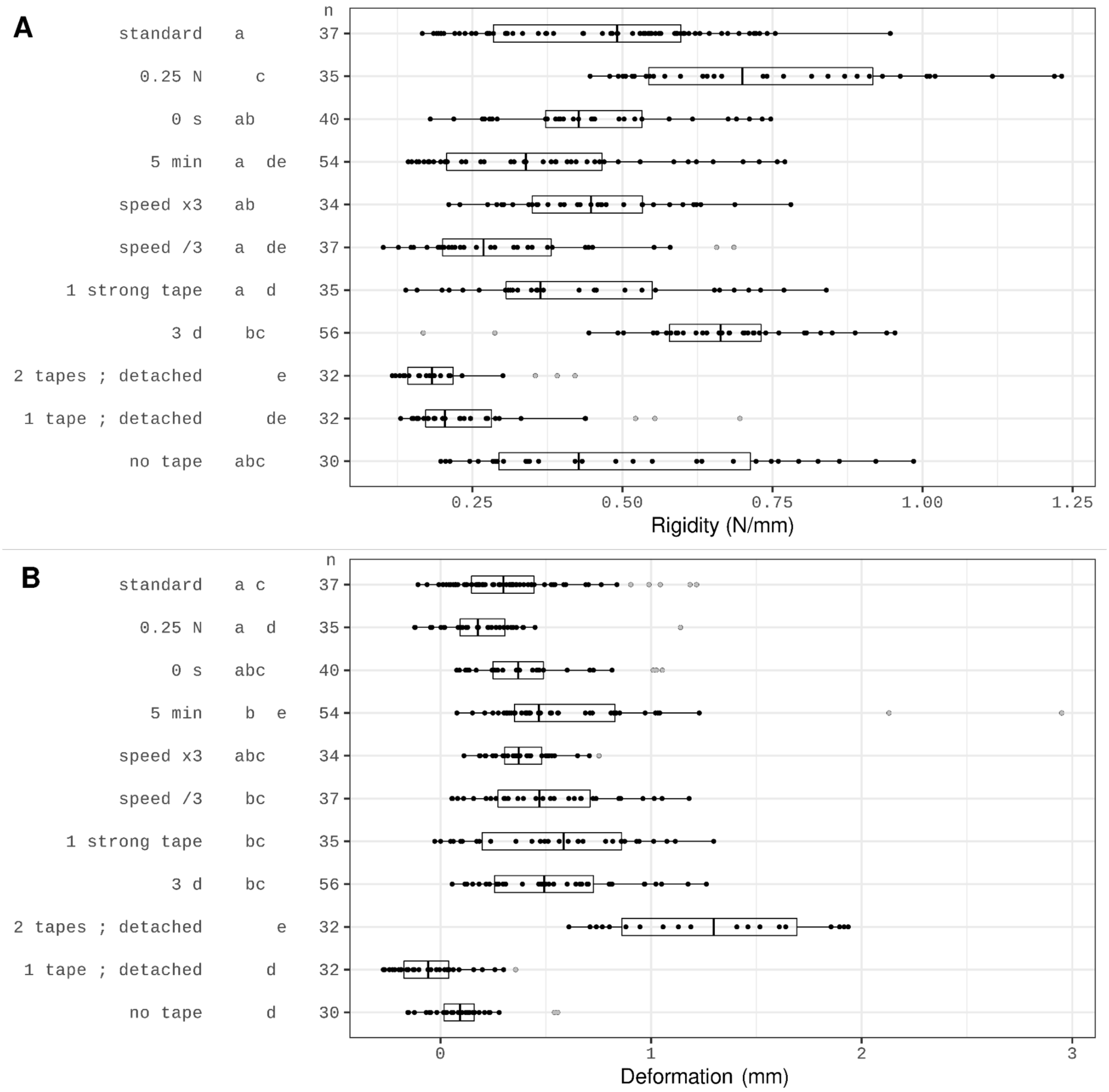


Figure S5. (A) Rigidity and (B) deformation for control adhesion assays with *D. melanogaster.* Same legend as Fig. S1. The various protocols are described in Table S2. Note that for the 0.25 N protocol the rigidity is calculated from the beginning to the end of the compression phase, and is thus overestimated due to the curve concavity visible on Fig. S4.

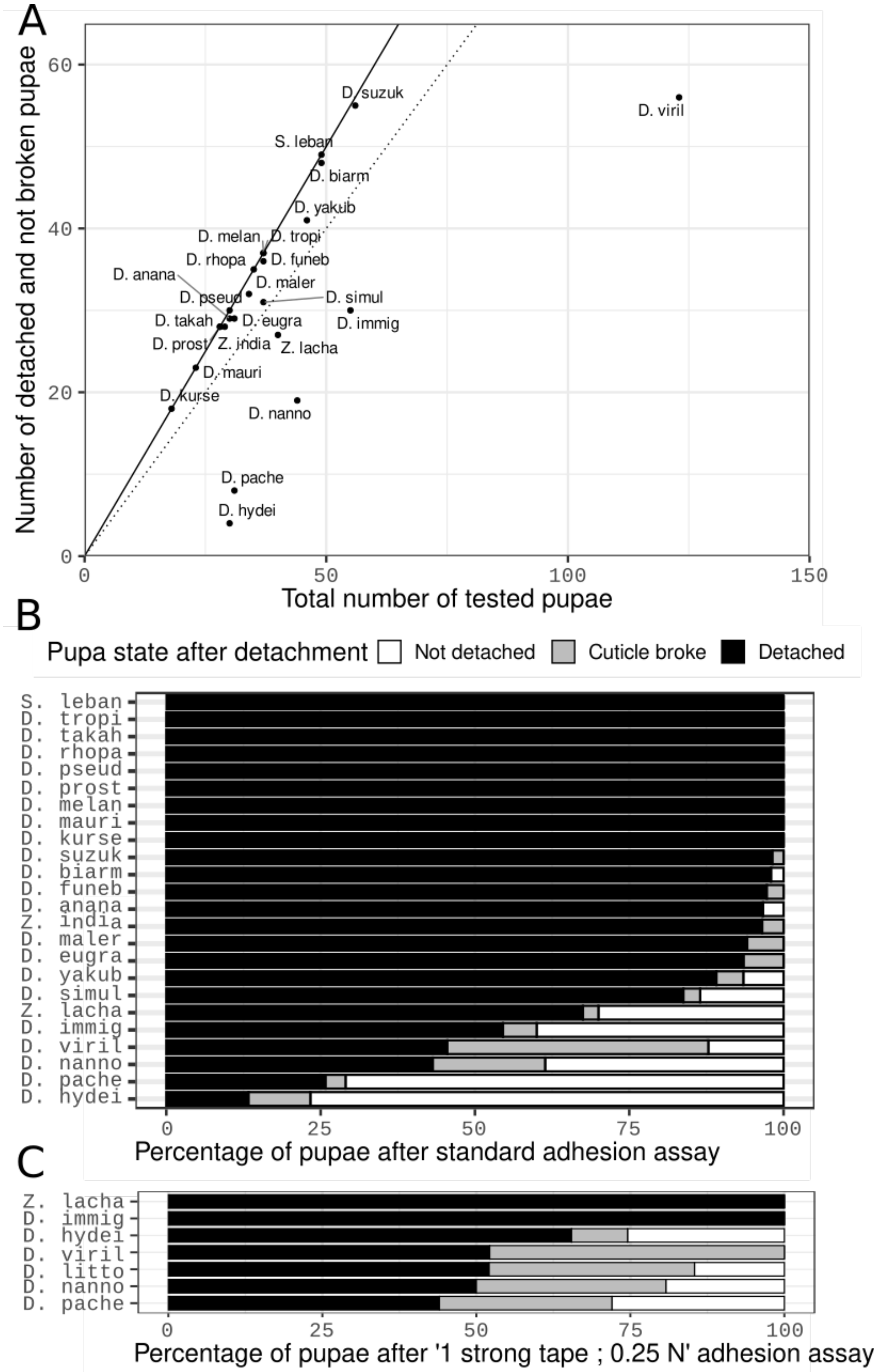


Figure S6. Number of pupae detached, not detached or broken during the standard and the ‘1 strong tape ; 0.25 N’ protocols. (A) Number of pupae detached and not broken over the total number of tested pupae with the standard protocol (see also Table S3). The black line is of slope 1 and the dotted line of slope 0.8. Species with less than 80% of detached pupae were tested further with the ‘1 strong tape ; 0.25 N’ protocol. (B, C) Percentage of detached, not detached and broken pupae obtained with the standard protocol (B) and with the ‘1 strong tape ; 0.25 N’ protocol (C). Species are ordered by increasing numbers of detached pupae. Species names are abbreviated using their first five letters. See Table S4 for numbers of pupae.

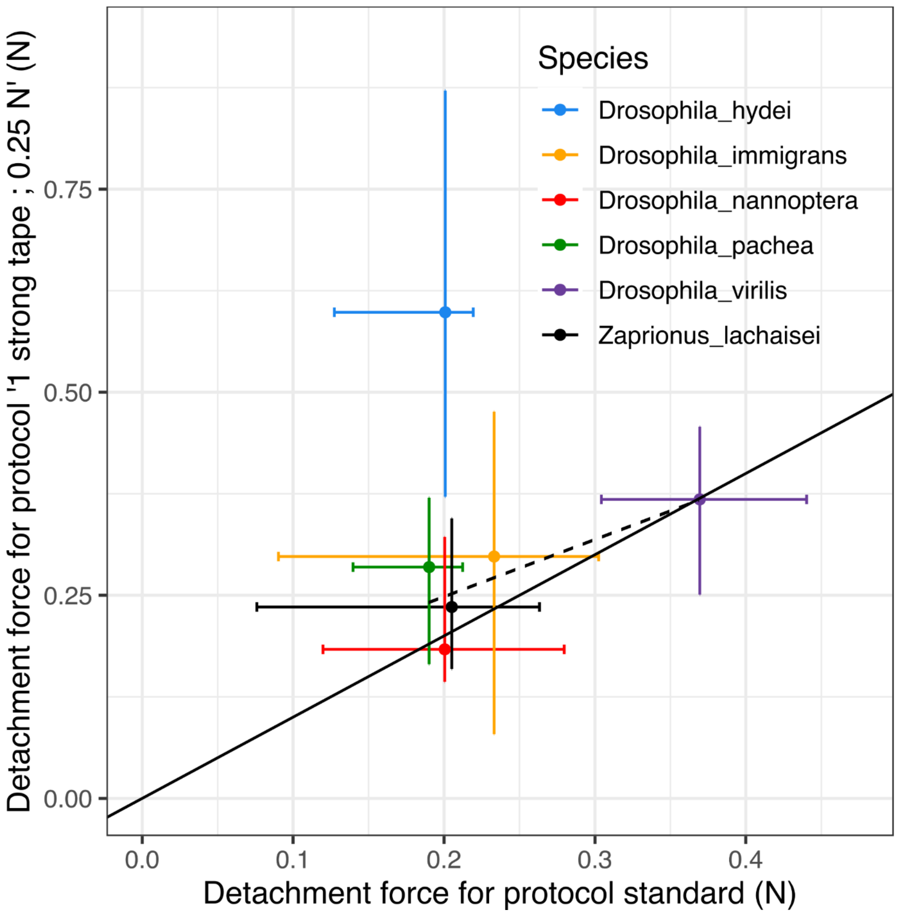


Figure S7. Effect of protocol on detachment force. Detachment force obtained with the '1 strong tape ; 0.25 N' protocol versus detachment force obtained with the standard protocol for six species indicated on the panel. The centre of the crosses correspond to the median. Horizontal and vertical bars represent the first and third quartiles. Pupae that are either detached, not detached or broken during adhesion assays are included. The solid black line represents a slope of 1 and the dotted line is the regression line between the median values of the five represented species (with *D. hydei* excluded). Regression: $r^2 = 0.82$, $p = 0.034$.

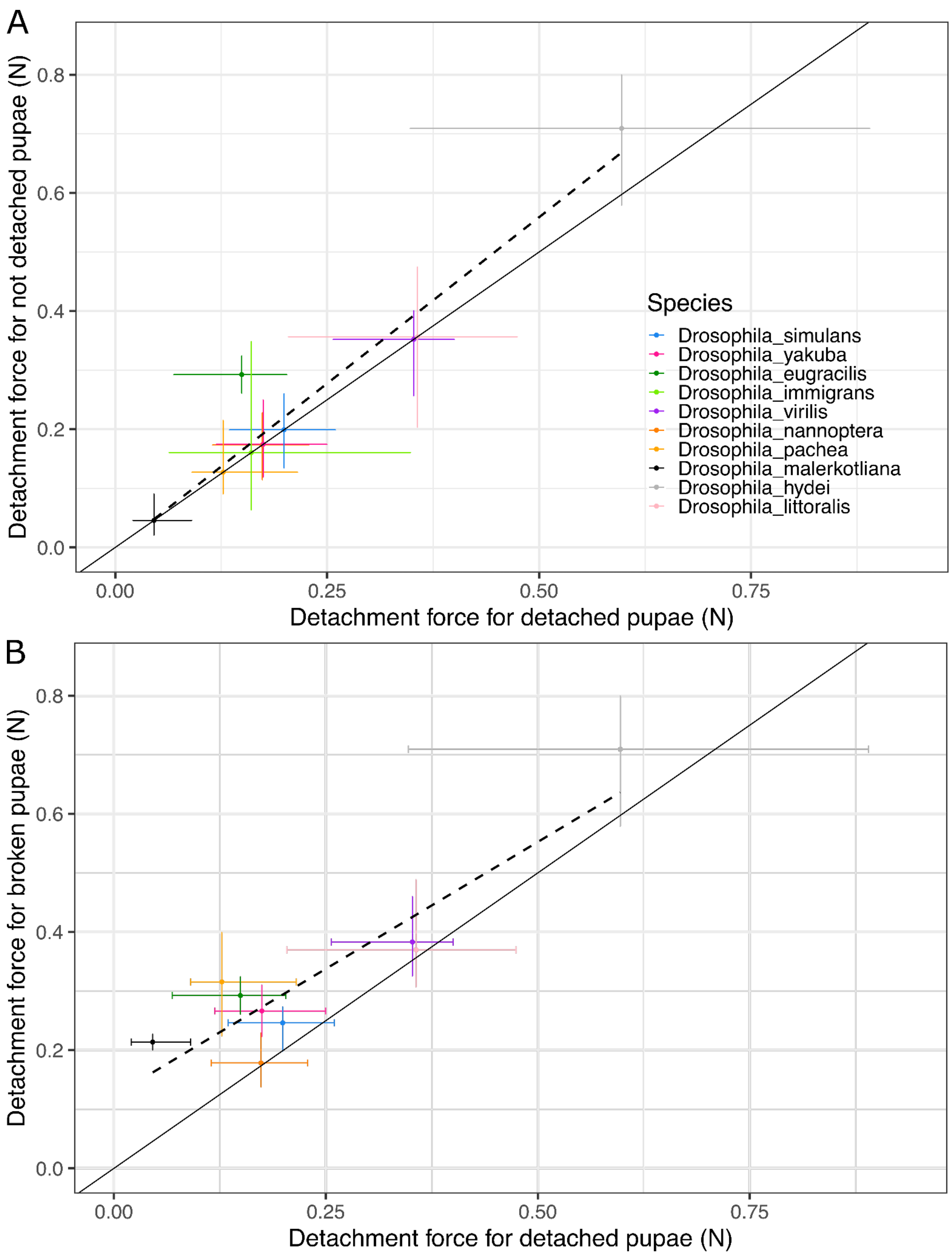


Figure S8. Detachment force of undetached or broken pupae compared with detachment force of detached pupae. Pupae detached with the standard or the "1 strong tape ; 0.25 N" protocol are used for the ten species that harboured at least two undetached or broken pupae, except for *D. hydei* where only values obtained with the "1 strong tape ; 0.25 N" protocol are shown. (A) Median detachment force of undetached pupae versus the median detachment force of detached pupae. (B) Median detachment force for broken pupae versus median detachment force for detached pupae for the 10 species that harboured at least 2 undetached or broken pupae. Regressions: A: $r^2 = 0.93$, $p = 8.2e\text{-}06$ ; B: $r^2 = 0.83$, $p = 6e\text{-}04$. Same legend as Figure S7.

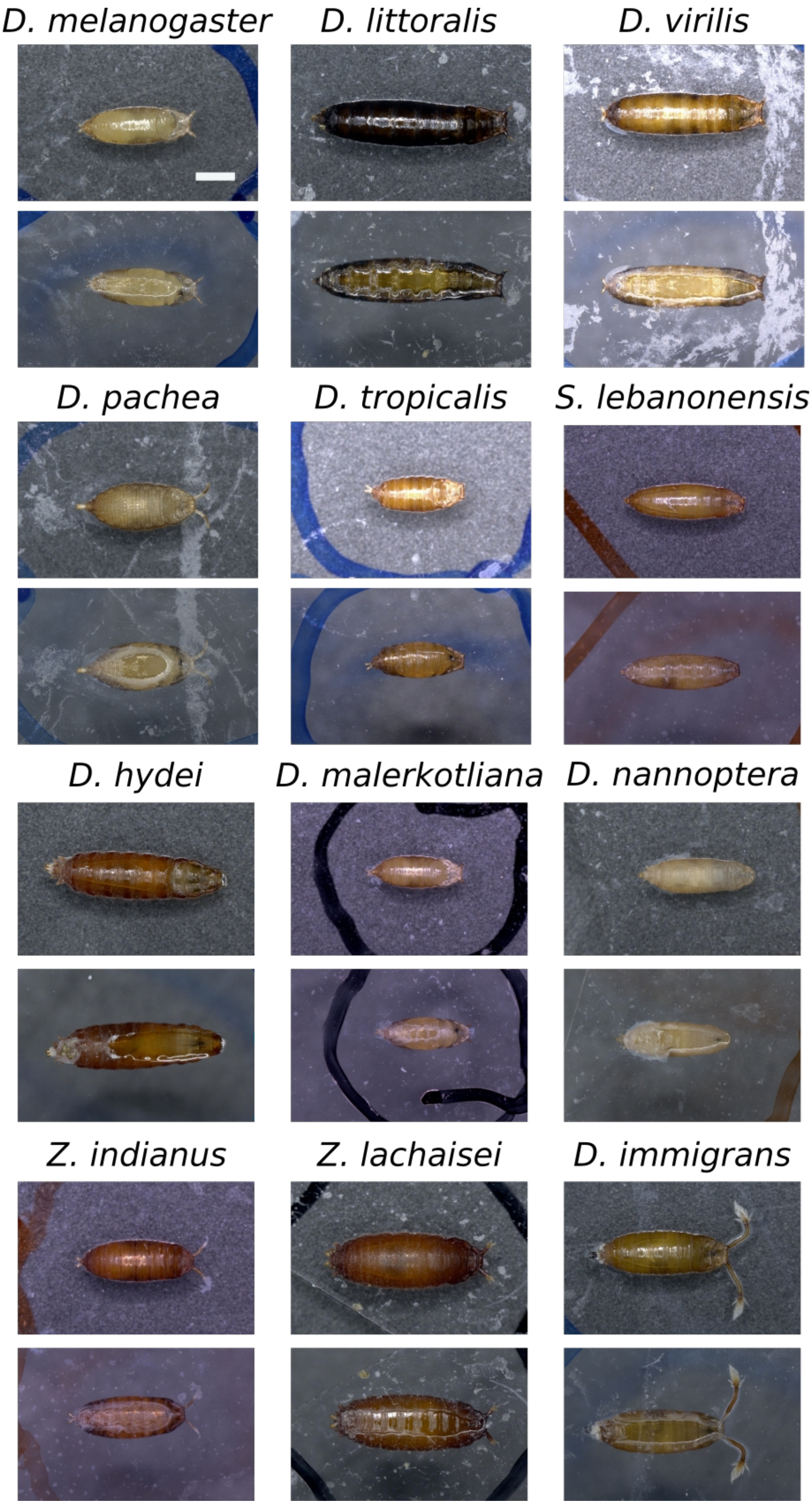


Figure S9. Representative examples of pupa dorsal and ventral pictures. Pupae from *D. melanogaster* and eleven species which display differences in size and shape (*D. littoralis, D. virilis, D. pachea, D. tropicalis, Scaptodrosophila lebanonensis, D. hydei, D. malerkotliana, D. nannoptera, Zaprionus indianus, Z. lachaisei* and *D. immigrans*). Wandering larvae were deposited on glass slides and pictures were taken 15-21 hours later. Marks of felt-pen are visible on slides. All pictures are of the same scale. Scale bar is 1 mm.

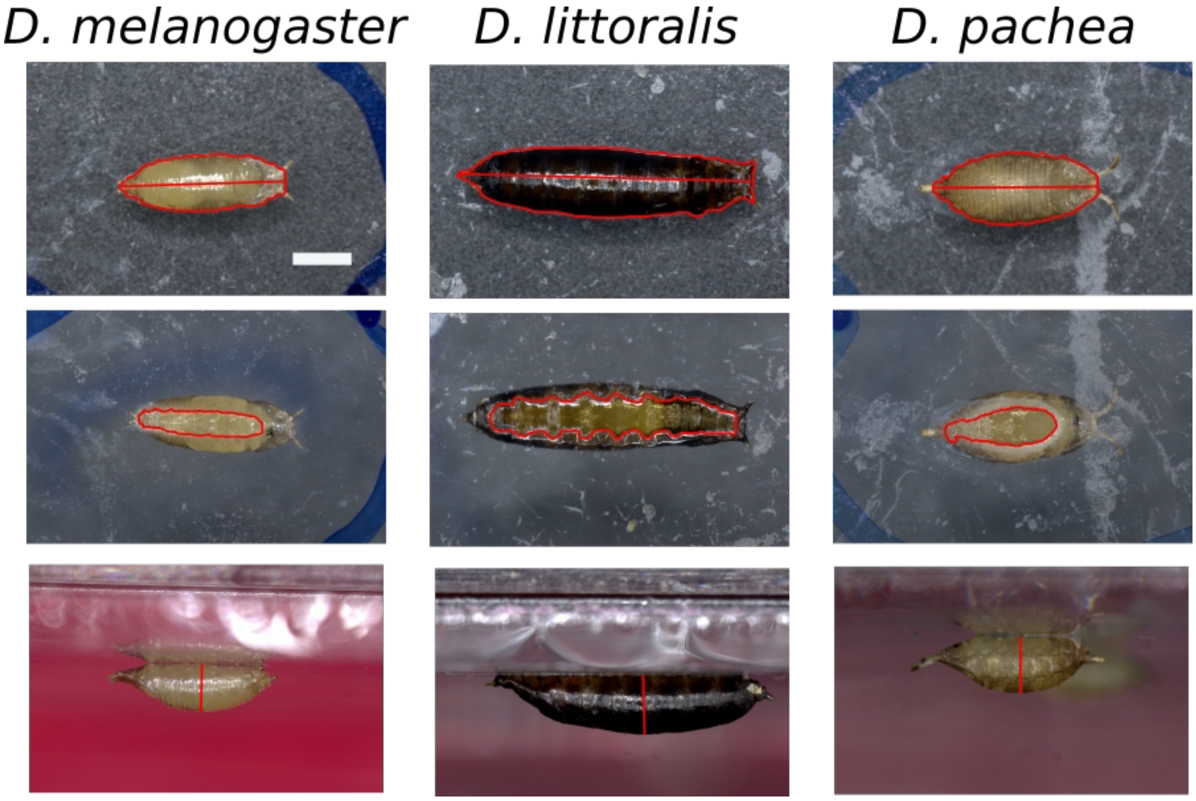


Figure S10. Measurements using pupa pictures. Pupal length, area, height and glue area for *D. melanogaster* and two species representative of differences in size and shape (*D. littoralis* and *D. pachea*) are shown. Pupal length and area are measured on dorsal pictures (first row), glue area is measured on ventral pictures (second row) and pupal height on profile pictures (third row). Outlines and lines used for measurements are shown in red. Same legend as Fig. S9.

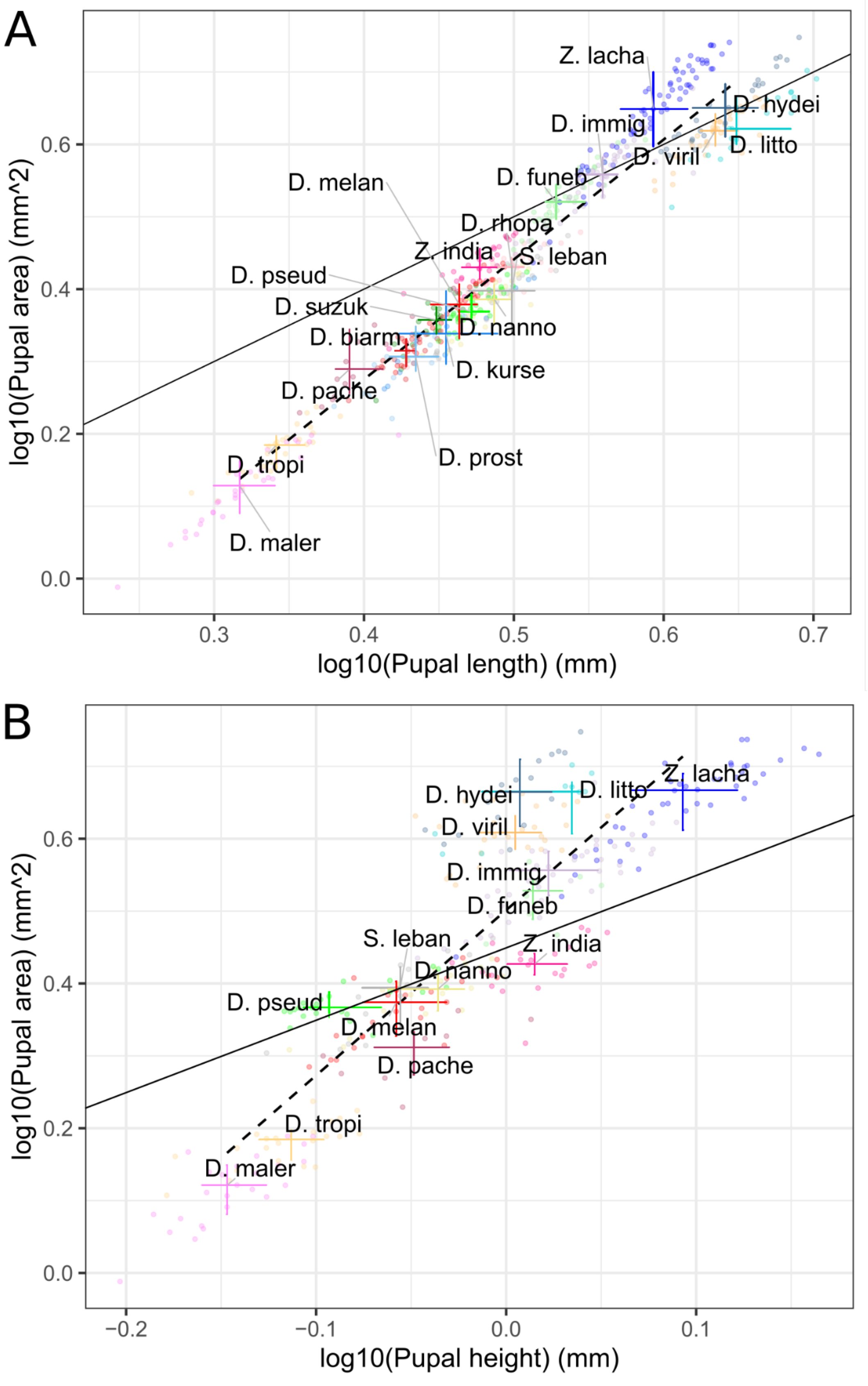


Figure S11. Pupal area versus pupal length and heightl. Species names are abbreviated using their first five letters and species are distinguished with colours. Each dot corresponds to a single pupa detached during adhesion assay. The centre of the crosses corresponds to the median. Horizontal and vertical bars represent the first and third quartiles. The solid black line represents a slope of 1. The dotted line is the regression line between the median values for each species. Regressions: A: $r^2 = 0.96$, $p < 2.2e\text{-}16$ ; B: $r^2 = 0.85$, $p < 2.2e\text{-}16$.

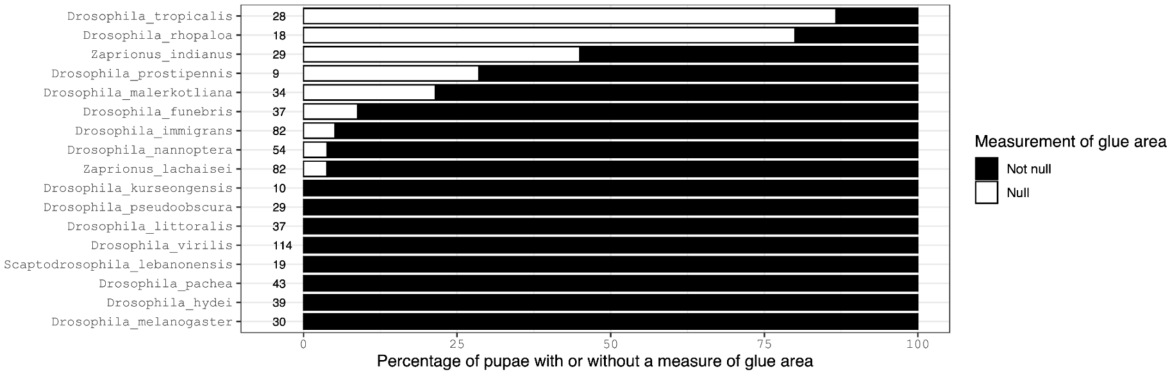


Figure S12. Percentage of pupae for which no glue area was detected on the ventral pictures See also Fig. S9. Same legend as Fig. S6B. For each species the total number of examined pupae is indicated.

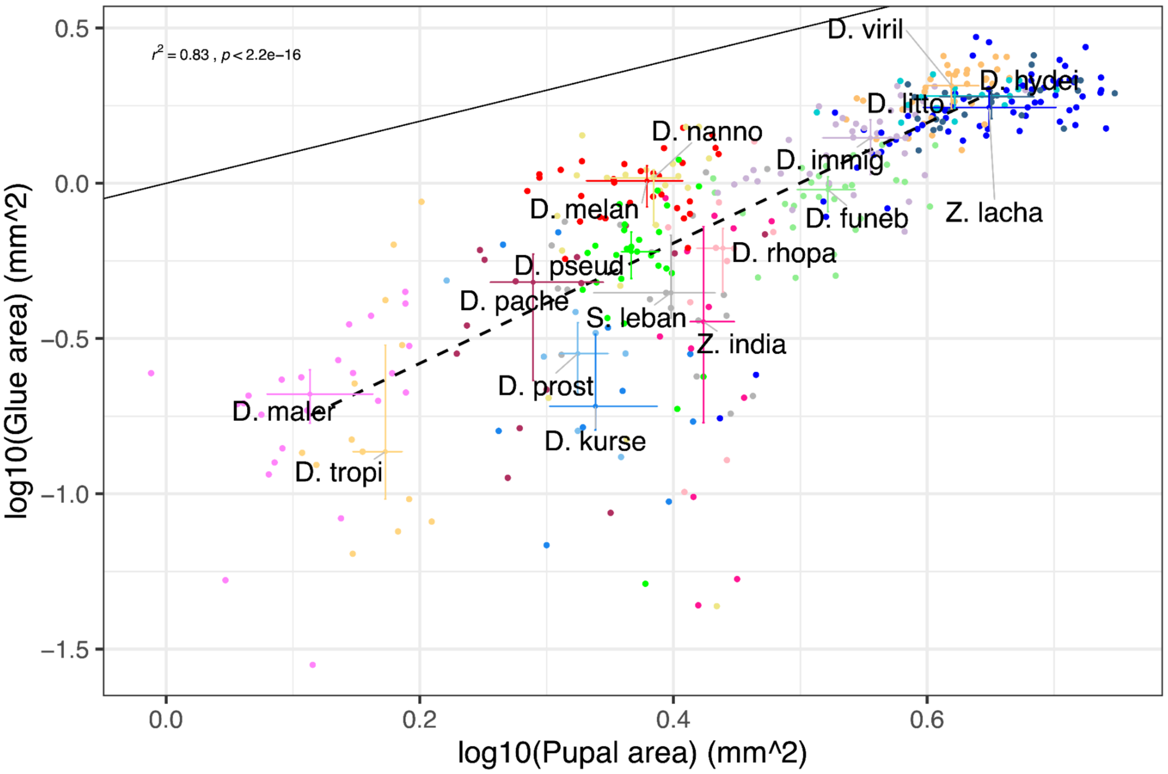


Figure S13. Glue area versus pupal area. Points where the glue area is null are not included. Regression: $r^2 = 0.83$, $p < 2.2e-16$. Same legend and colours as Fig. S11.

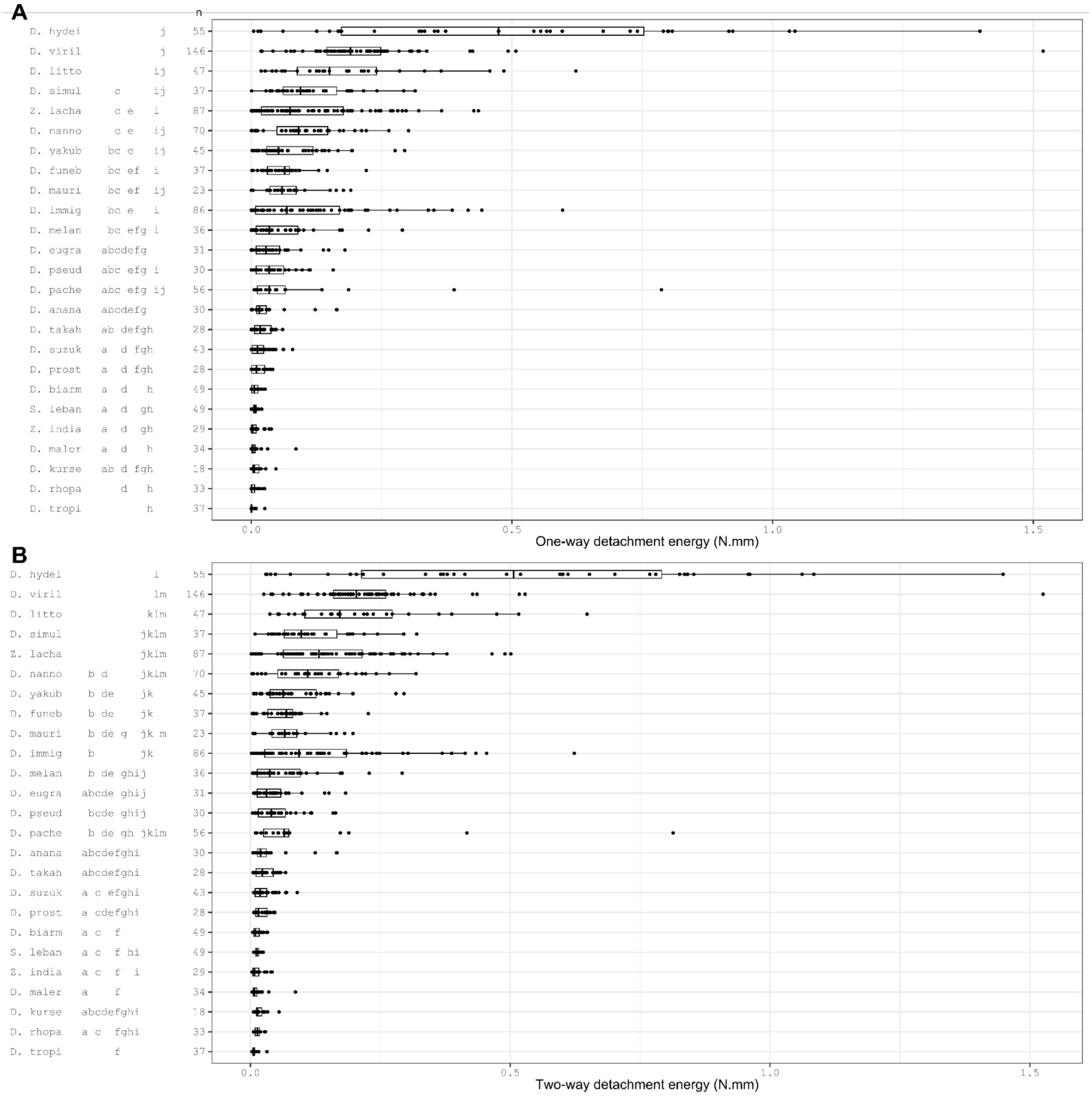


Figure S14. One-way and two-way detachment energy for 25 Drosophila species. Each dot corresponds to a single pupa detached with the standard or the "1 strong tape ; 0.25 N" protocol. For *D. hydei,* only values obtained with the "1 strong tape ; 0.25 N" protocol are shown. Values below '*n*' indicate the total number of pupae tested for each species. Species are ordered by increasing median detachment force, with *D. hydei* and *D. tropicalis* having the highest and lowest median detachment forces, respectively. Same legend as Fig. S1.

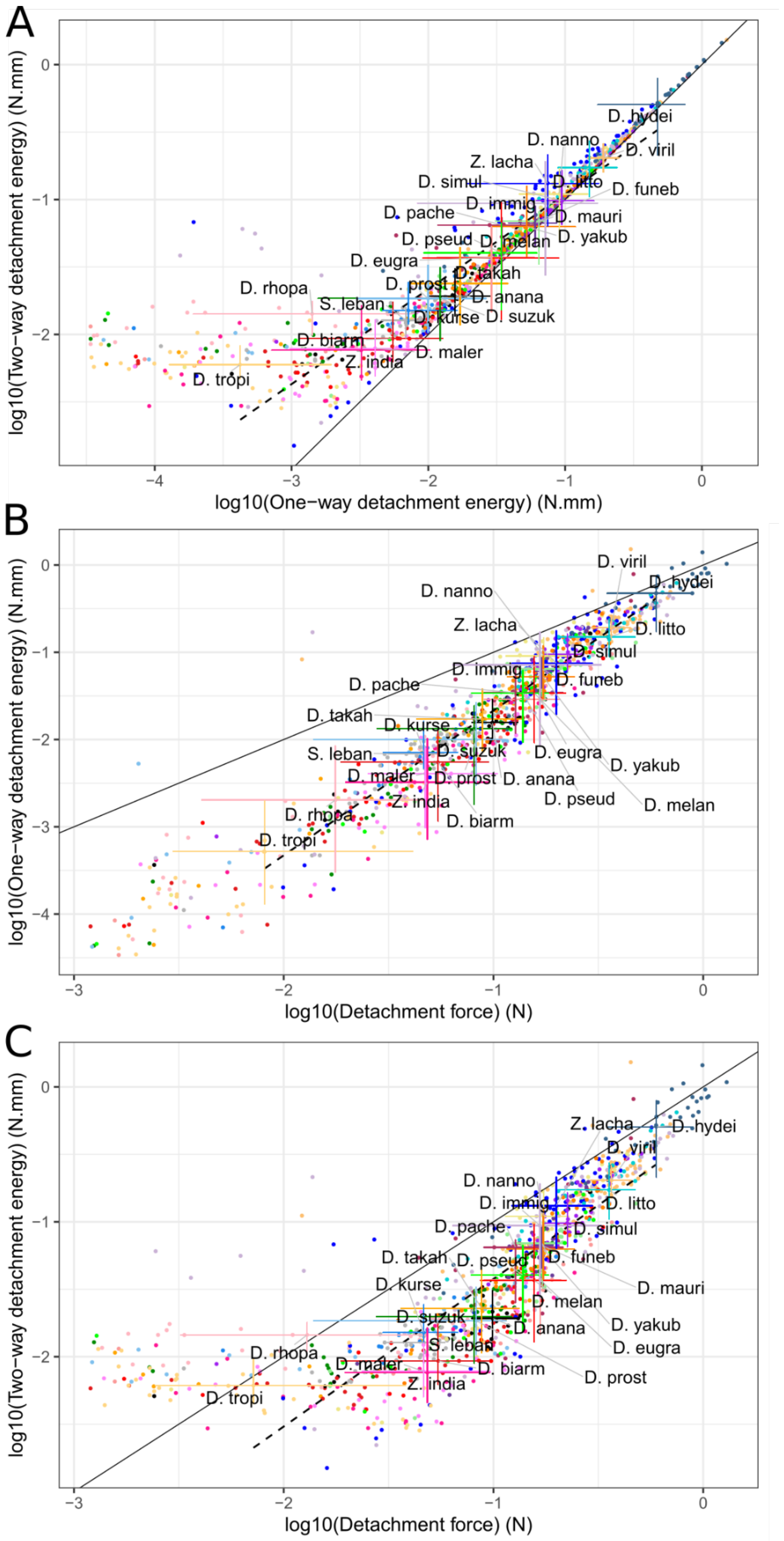


Figure S15. Detachment force and energies for 25 Drosophila species. (A) Two-way detachment energy versus one-way detachment energy, (B) one-way detachment energy versus detachment force, (C) two-way detachment energy versus detachment force. Regions under 0.03 N are not displayed. Regions under 0.03 N are not used in the regression. A:

$r^2 = 0.91$, $p < 2.2e\text{-}16$ ; B: $r^2 = 0.96$, $p < 2.2e\text{-}16$ ; C: $r^2 = 0.8$, $p < 2.2e\text{-}16$. Same legend and colours as Fig. S11.

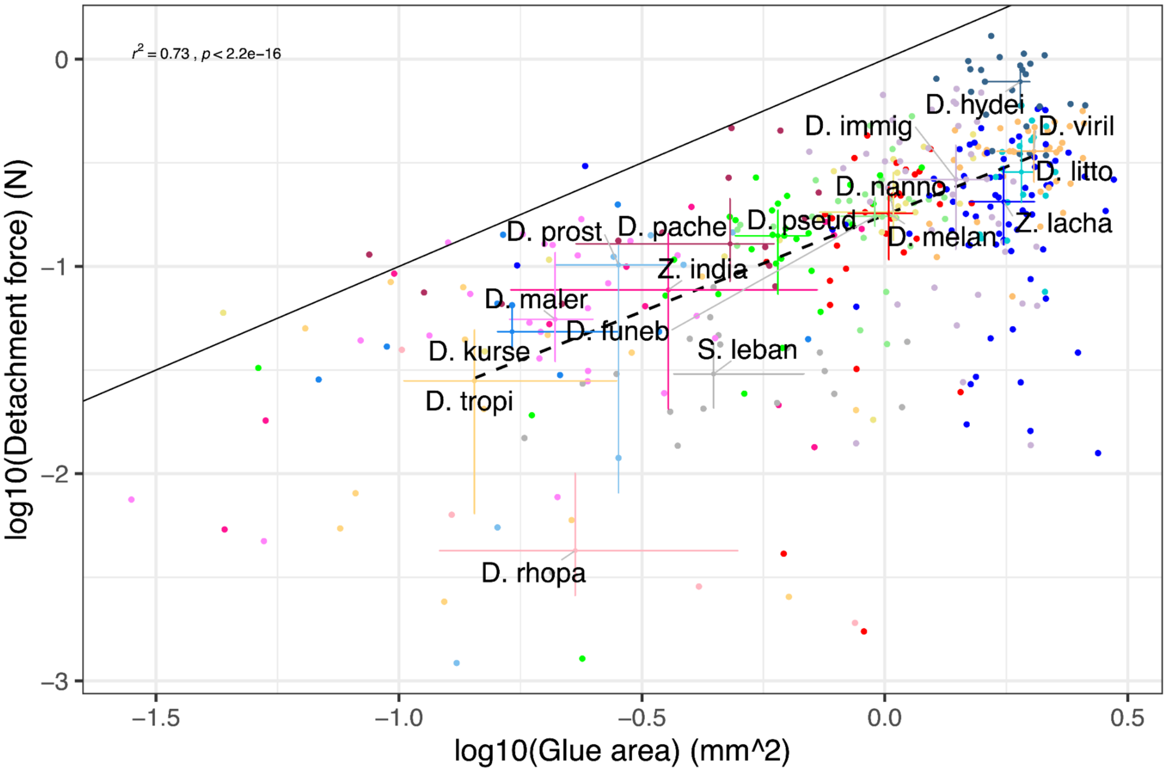


Figure S16. Glue area versus detachment force. Points where the glue area is null are not included. Regression: $r^2 = 0.73$, $p < 2.2e\text{-}16$. Same legend and colours as Fig. S11.

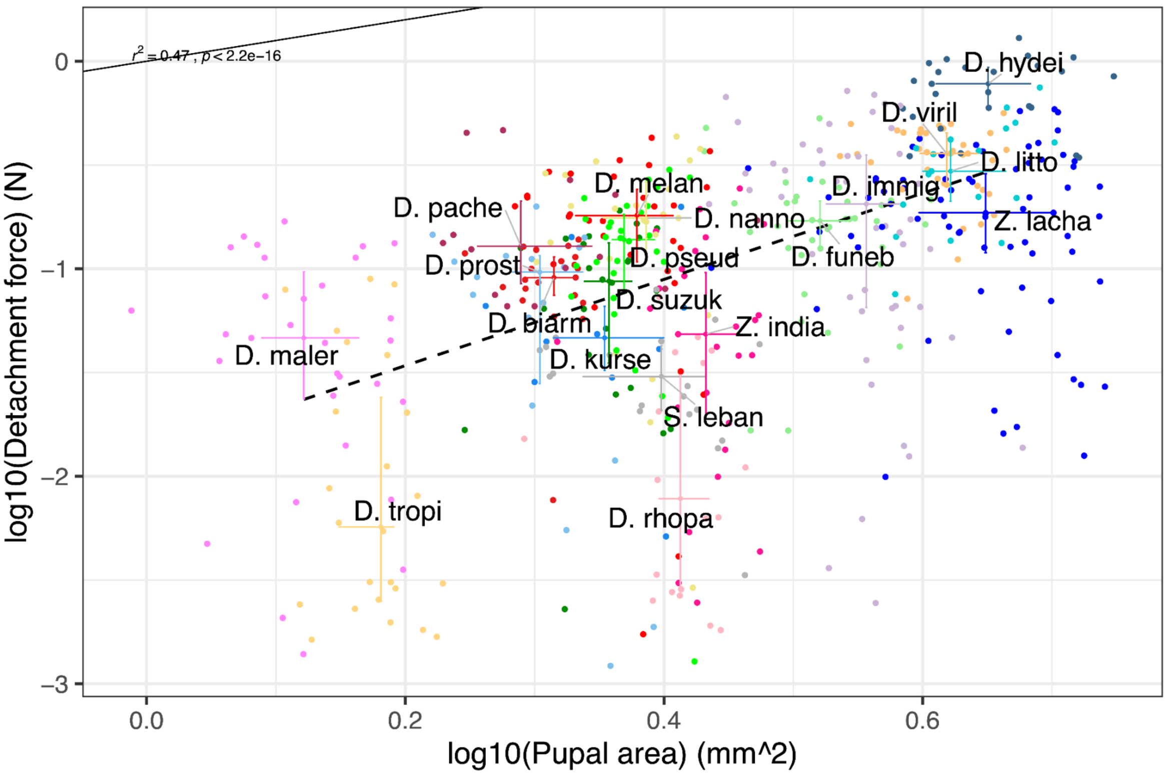


Figure S17. Detachment force versus pupal area. Regression coefficient $r^2 = 0.47$, $p < 2.2e\text{-}16$. Same legend and colours as Fig. S11.

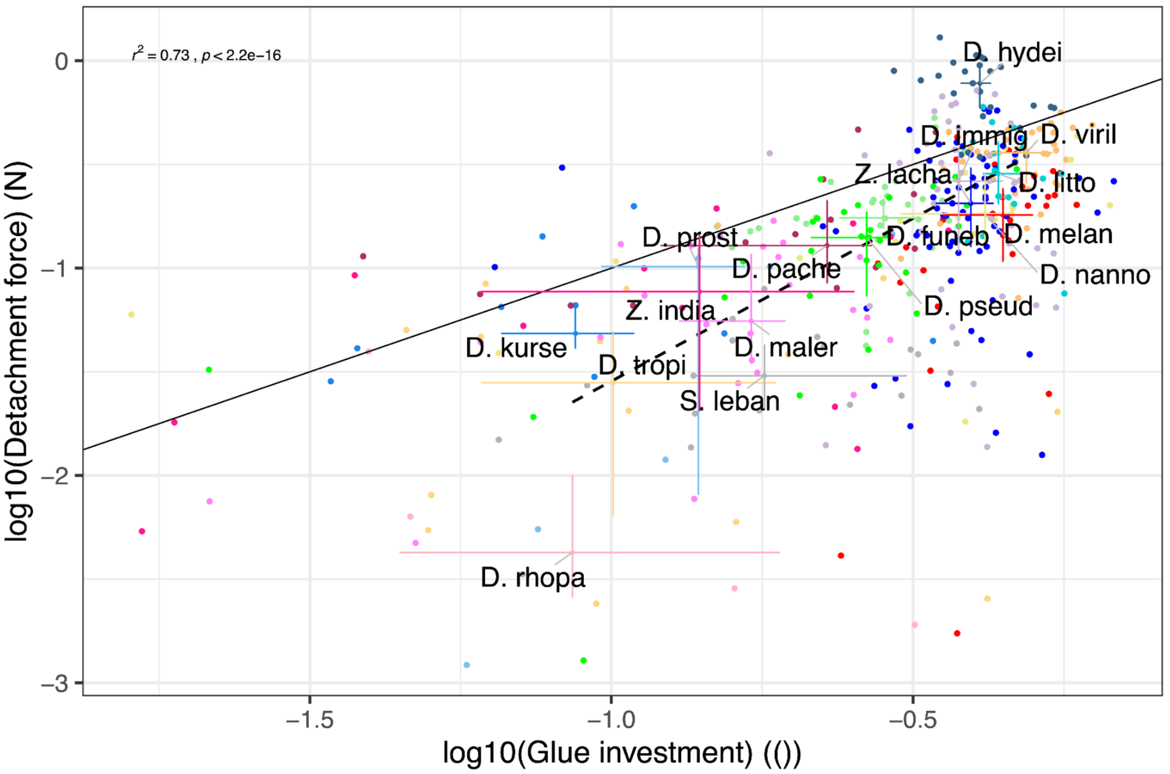


Figure S18. Detachment force versus glue investment. Regression coefficient $r^2 = 0.73$, $p < 2.2e\text{-}16$. Same legend and colours as Fig. S11.

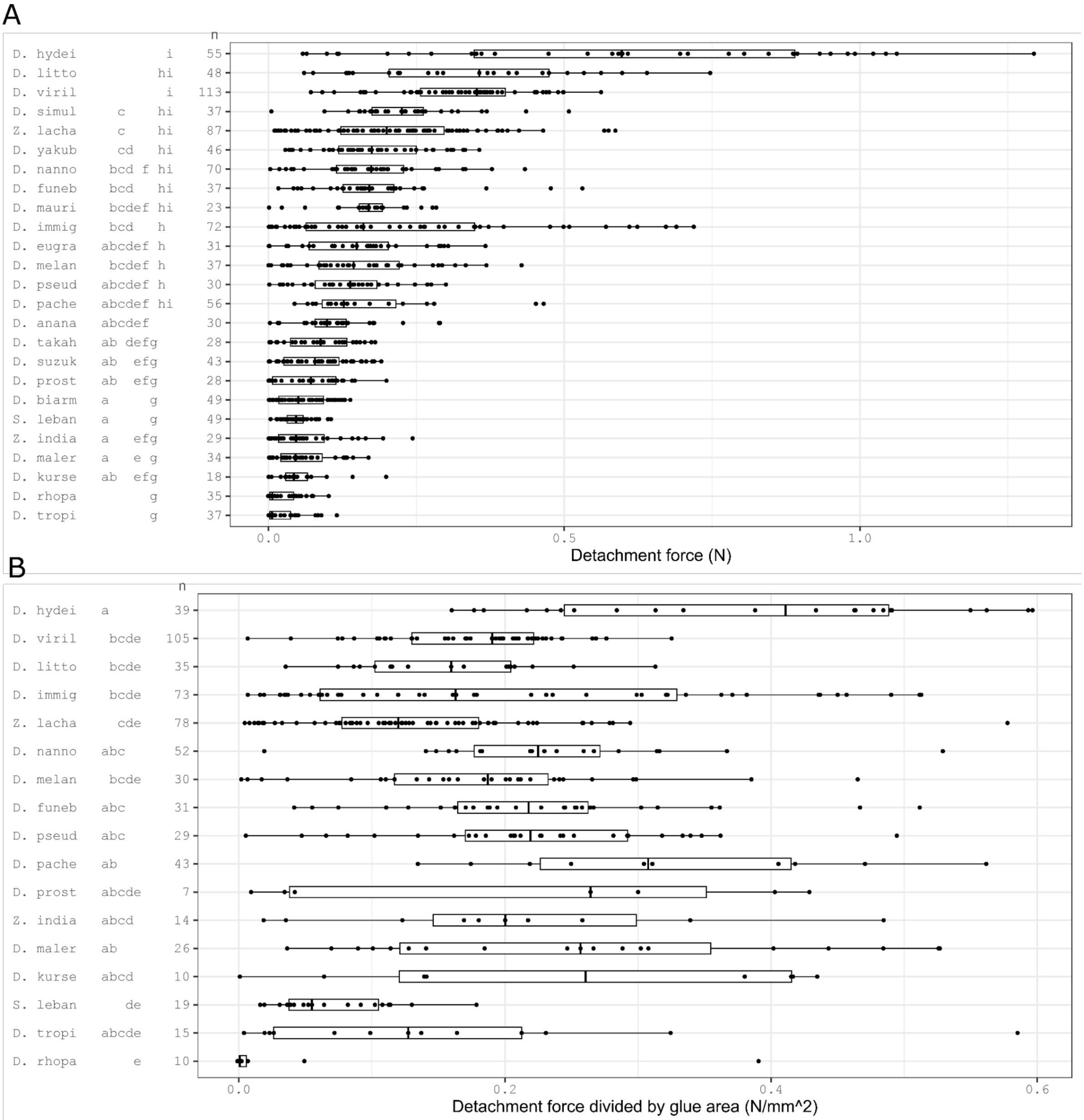


Figure S19. Detachment force and strength for several Drosophila species. Each dot corresponds to a single pupa tested with standard or "1 strong tape ; 0.25 N" protocol. For *D. hydei* only data points obtained with the "1 strong tape ; 0.25 N" protocol are shown. Only measures of detached pupae are shown here. See Fig. S20 for broken and undetached pupae. Same legend as Fig. S1.

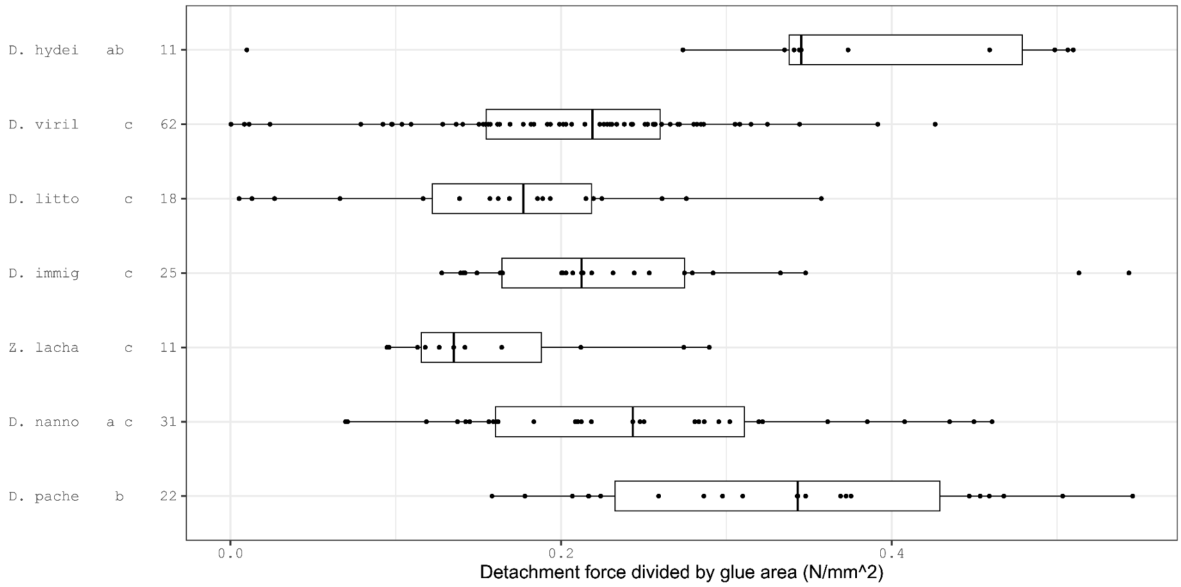


Figure S20. Detachment force per glue area for pupae that were not detached or broken during adhesion assays. Species with less than two pupae being not detached or broken are not displayed. For reasons of legibility the highest data points resulting from division by extremely small numbers (representing 5% of all data) are not shown. For *D. hydei* only data points obtained with the "1 strong tape ; 0.25 N" protocol are shown. Same legend as Fig. S1.

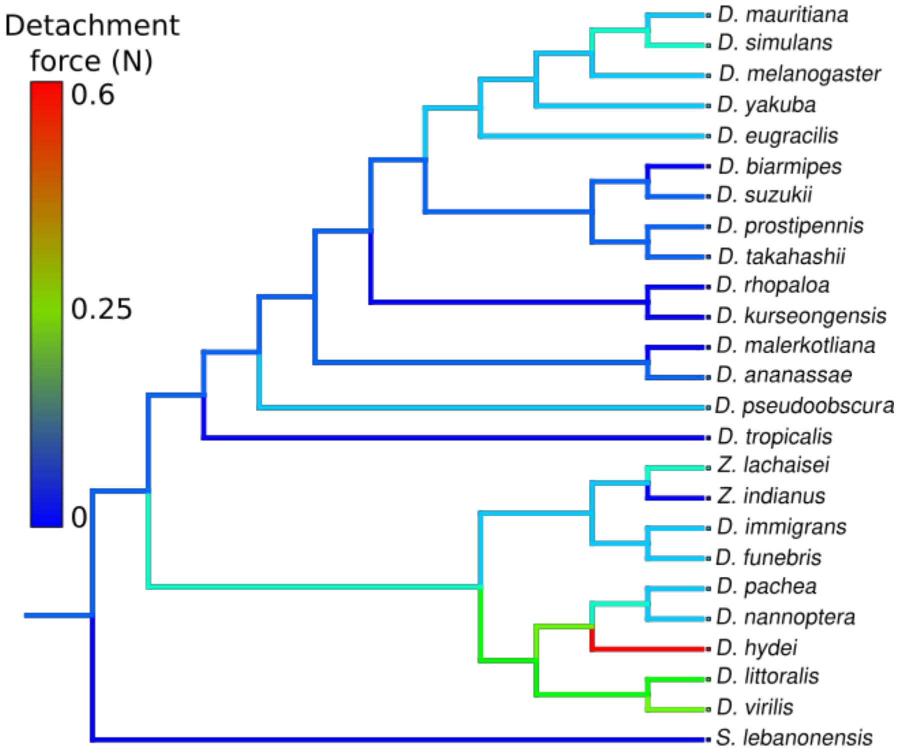


Figure S21. Evolution of the adhesion force across the phylogeny. Species branches are coloured according to their median detachment force, from blue to red, respectively, corresponding to low and high detachment force. Ancestral branches are coloured based on a parsimony algorithm. For example, *Z. indianus* is not adhesive, its branch is coloured in dark blue and *Z. lachaisei* has a medium adhesion, its branch is coloured in green. The most parsimonious scenario is that the common ancestral branch for both species has a low adhesion, its branch is coloured in light blue. Phylogenetic tree is adapted from (Suvorov et al., 2022) and tree branch lengths do not represent real distances.

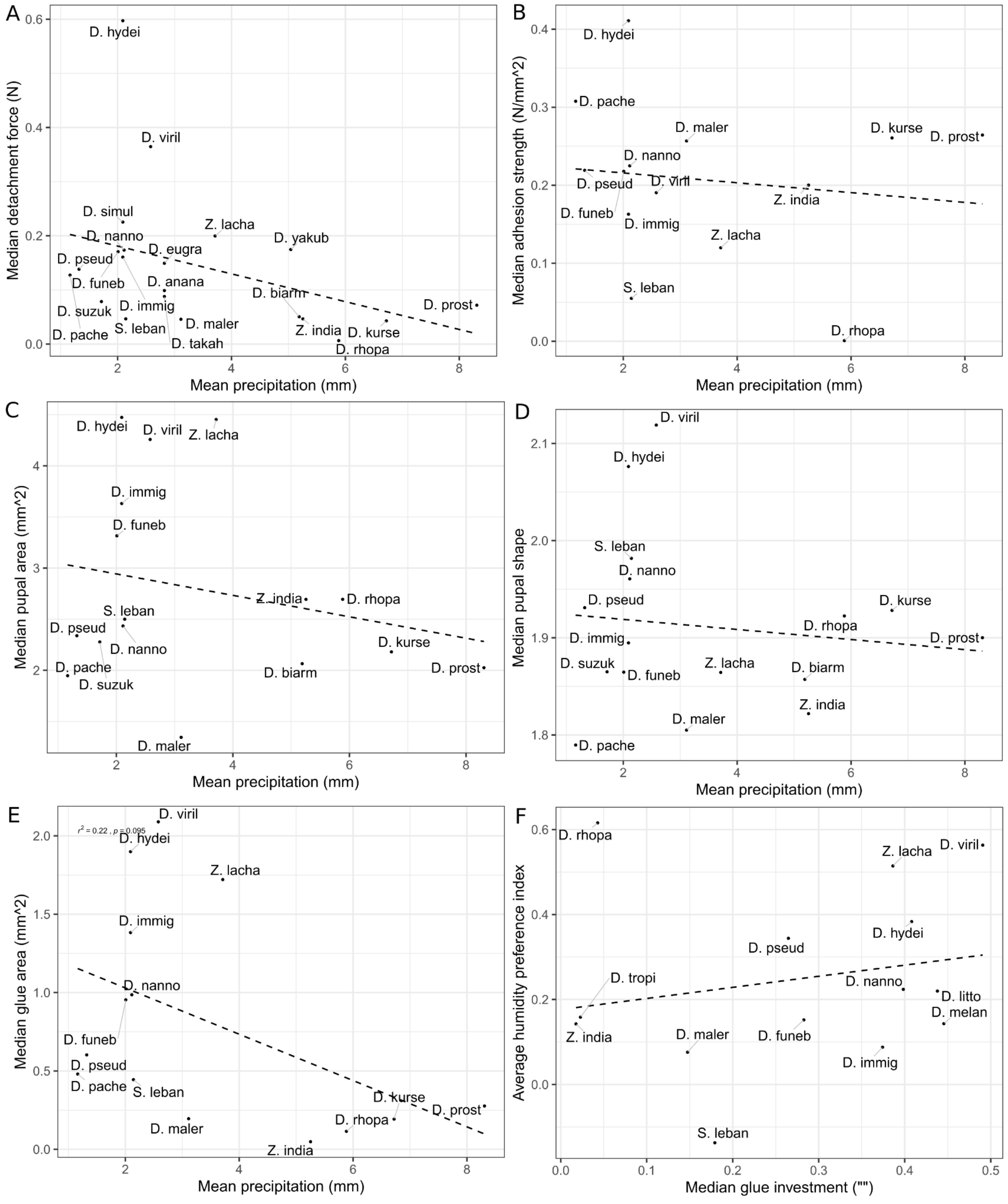


Figure S22. Geometrical and mechanical properties (Detachment force, adhesion strength, pupal area, pupal shape and glue area) versus mean precipitation and Average humidity preference index versus glue investment. (A) Median detachment force (N) versus mean precipitation (mm). (B) Median adhesion strength (N/mm²) versus mean precipitation (mm). (C) Median pupal area (mm²) versus mean precipitation (mm). (D) Median pupal shape versus mean precipitation (mm). (E) Median glue area (mm²) versus mean precipitation (mm). (F) Average humidity preference index versus median glue investment. Linear regression: A: $r^2 = 0.14$, $p = 0.092$ ; B: $r^2 = 0.0018$, $p = 0.64$; C: $r^2 = 0.056$, $p = 0.038$; D: $r^2 = 0.15$, $p = 0.065$; E: $r^2 = 0.022$, $p = 0.095$; F: $r^2 = 0.043$, $p = 0.48$;.

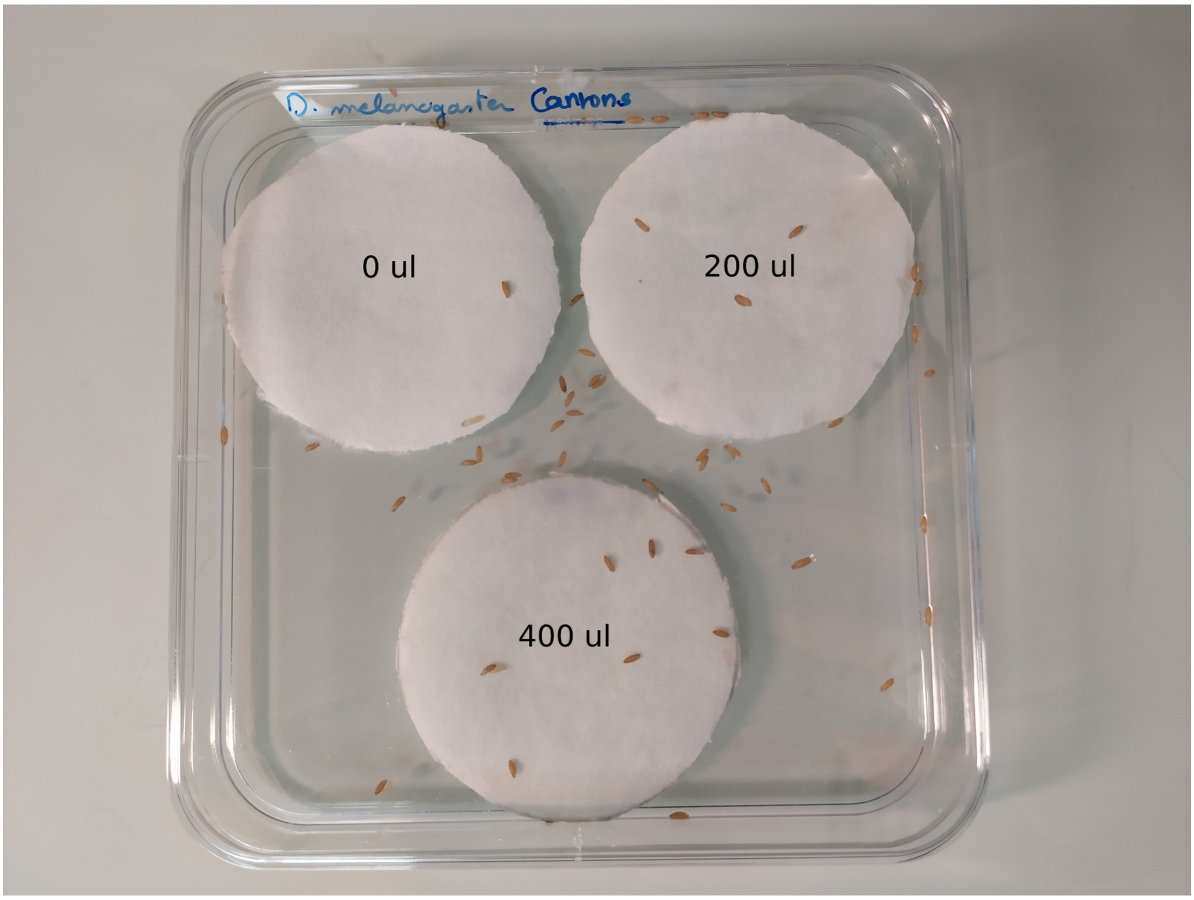


Figure S23. Representative picture of our humidity preference experiment with *D. melanogaster.* In a 12 cm x 12 cm square Petri dish, three round pieces of Whatman paper are attached with double sided tape. The first paper circle was left dry, the second wetted with 200 µL of fresh tap water and the third with 400 µL. *D. melanogaster* L3 wandering larvae were deposited at the centre of the box and allowed to choose a site for pupariation. In this picture, a large proportion of animals ended up on the plastic. The average humidity index for *D. melanogaster* is 0.14.

### Supplementary Tables

**Table S1. List of Diptera species used in this study, and available details regarding collection dates and places.**

| Species | Stock | Origin | Temperature of culture (°C) |
|---|---|---|---|
| *Drosophila melanogaster* | Canton S | Given by R. Karess. | 25 |
| *Drosophila eugracilis* | | Given by N. Gompel. Isofemale line #11, India, Karnataka, Bengaloru, GKVK campus, Area E10 (920 m.) [13° 4'14.60"N 77°35'7.01"E], 1 Jan 2013, B Prud'homme/N Gompel legacy. | 22 |
| *Drosophila biarmipes* | G224 | Given by N. Gompel. Line G224 in lab collection (361.0-iso 1-11, Genome strain 1, derived from San Diego Stock Center line: 14023-0361.00. Cambodia, Ari Ksatr. | 22 |
| *Drosophila takahashii* | San Diego Stock Center 14022-0311.07 Isofemale #01 and #03 | Given by N. Gompel. Isofemale line #01 and #03: India, Karnataka, Bengaloru, GKVK campus, Area E10 (920 m.) [13° 4'14,60"N 77°35'7.01"E], 1 Jan 2013, B Prud'homme/N Gompel legacy | 22 |
| *Drosophila rhopaloa* | BaVi067 | Given by N. Gompel. Line BaVi067 from Vietnam, Hanoi Ba Vì, near Vân Hòa [21°04′N, 105°22′E], March 2005, H. Takamori legacy. | 22 |
| *Drosophila prostipennis* | San Diego Stock Center 14022-0291.00 | Given by N. Gompel. San Diego Stock Center 14022-0291.00, Taiwan, Wulai, 04/06/1968 (L. Throckmorton legacy.) | 22 |
| *Drosophila ananassae* | Isofemale #01 | Given by N. Gompel. Isofemale #01: India, Karnataka, Bengaloru, GKVK campus, Area E10 (920 m.) [13° 4'14,60"N 77°35'7.01"E], 1 Jan 2013, B Prud'homme/N Gompel legacy. | 22 |
| *Drosophila kurseongensis* | SaPa058 | Given by N. Gompel. Line SaPa058 from Vietnam, Lao Cai, Sa Pa [22°20′N 103°52′E], March 2009, H. Takamori leg (gift from T. Matsuo) | 22 |
| *Drosophila funebris* | | From 10 larvae approximately collected by M. Monier in Pennautier, France in July 2021. | 22 |
| *Drosophila pachea* | 15090-1698.01_14.2 | Inbred line of 15090-1698.01 (11 generations of single pair crosses) established in 1997 from individuals caught in Arizona, USA. Given by M. | 25 |

| | | | |
|---|---|---|---|
| | | Lang. | |
| *Drosophila nannoptera* | 15090-1692.00 | Cornell Drosophila Stock Center #15090-1692.00. Established in 1992 from individuals caught in Oaxaca, Mexico. Given by M. Lang. | 25 |
| *Scaptodrosophila lebanonensis* | | Given by S Prigent – collected by J. David in Prunay, France | 25 |
| *Drosophila pseudoobscura* | 14011-0121.94 | Cornell Drosophila Stock Center #14011-0121.94. Origin: Anderson, Mesa Verde, Colorado, USA. | 25 |
| *Drosophila hydei* | Vincennes | Vincennes park (Paris, France). Collected by S. Prigent in 2020 (13). | 22 |
| *Drosophila immigrans* | Vincennes | Vincennes park (Paris, France). Collected by S. Prigent in 2020 (13). | 22 |
| *Drosophila virilis* | 15010-1051.86 | Cornell Drosophila Stock Center #15010-1051.86. Given by A. Farlow from C. Schloetterer's lab in 2010. Origin: Texmelucan, Puebla, Mexico. | 22 |
| *Drosophila littoralis* | | Given by S. Prigent. Collected in Rostov (Russia) by J. David. Unknown date. | 25 |
| *Drosophila malerkotliana* | | Given by S. Prigent. Collected in Mayotte by J. David, A. Suwalski and A. Yassin in 2017. | 25 |
| *Drosophila mauritiana* | mau12 | Given by P. Andolfatto in May 2020. | 22 |
| *Drosophila simulans* | Vincennes | Vincennes park (Paris, France). Collected by S. Prigent in 2020 (13). | 22 |
| *Drosophila suzukii* | WT3 | Watsonville (USA). | 25 |
| *Drosophila tropicalis* | | Origin unknown. Given by P. Gibert. | 22 |
| *Drosophila yakuba* | Tai18E2 | Cornell Drosophila Stock Center #14021-0261.01. Tai forest, border of Liberia and Ivory Coast, 1983 given by P. Andolfatto in May 2020. | 22 |
| *Zaprionus lachaisei* | | Given by S. Prigent. Collected in Tanzania: East Usambara Mt, Amani (900 m) by D. Lachaise in 2002 (29). | 25 |
| *Zaprionus indianus* | | Given by S. Prigent. Collected in Grande Comore by J. David, A. Suwalski and A. Yassin in 2017. | 25 |

**Table S2. Adhesion assay protocols used. Rows indicate the different protocols used and columns the protocol parameters. Parameters similar to the standard protocol are represented by a hyphen -. Arrows pointing up or down, respectively, correspond to the sensor going up or down.**
1 strong tape: Gergonne acrylic double sided tape n° 01 2973X0X 22, 9 mm double sided tape was used.
Detached: before adhesion assays, pupae naturally attached on glass slides were manually detached with a paintbrush. Pupae damaged during this manual detachment were not kept. Detached pupae were then placed on a clean glass slide for adhesion measurement, with their ventral side in contact with the slide.
2 tapes: Manually detached pupae were placed on a piece of Tesa double-sided tape on a glass slide, with their ventral side in contact with the tape.
No tape: no tape was placed on the force sensor.

| | **Speed** | **Pause** | **Maximal force** | **Pause force** | **Type of tape** | **Presence of glue** | **Tape on the sensor** | **Pupa age** |
|---|---|---|---|---|---|---|---|---|
| **standard** | ↓0.016 mm/s ↑0.2 mm/s | 10 s | 0.07 N | 0.03 N | Tesa | Yes | Yes | 18 hours after larva deposition |
| **0 s** | - | 0 s | - | - | - | - | - | - |
| **5 min** | - | 300 s | - | - | - | - | - | - |
| **speed /3** | ↓0.33 mm/min ↑0.06 mm/s | - | - | - | - | - | - | - |
| **speed x3** | ↓0.0055 mm/s ↑0.6 mm/s | - | - | - | - | - | - | - |
| **0.25 N** | - | - | 0.25 N | 0.21 N | - | - | - | - |
| **1 strong tape** | - | - | - | - | Gergonne | - | - | - |
| **1 strong tape ; 0.25 N** | - | - | 0.25 N | 0.21 N | Gergonne | - | - | - |
| **1 tape ; detached** | - | - | - | - | - | Pupa detached | - | - |
| **2 tapes ; detached** | - | - | - | - | - | Pupa detached and placed on double sided Tesa tape | | |
| **no tape** | - | - | - | - | - | - | No | - |
| **3 d** | - | - | - | - | - | - | - | 3 days after larva deposition |

**Table S3. Number of pupae detached, not detached or broken during different adhesion protocols for *D. melanogaster.*** Rows indicate the protocols and columns indicate the number of pupae detached, not detached or broken during the given protocol.

| Protocol | Detached | Not detached | Cuticle broke |
|---|---|---|---|
| **0 s** | 29 | 0 | 11 |
| **0.25 N** | 32 | 3 | 0 |
| **1 strong tape** | 30 | 5 | 0 |
| **1 tape ; detached** | 31 | 1 | 0 |
| **2 tapes ; detached** | 20 | 6 | 6 |
| **3 d** | 36 | 4 | 16 |
| **5 min** | 43 | 3 | 8 |
| **no tape** | 30 | 0 | 0 |
| **speed /3** | 30 | 4 | 3 |
| **speed x3** | 33 | 1 | 0 |
| **standard** | 37 | 0 | 0 |

**Table S4. Number of pupae detached, not detached or broken during the '1 strong tape ; 0.25 N' for 6 species and the standard protocols for 24 species other than *D. melanogaster.*** Rows indicate the species and columns indicate the number of pupae detached, not detached or broken during the given protocol. Sub-totals and totals are in boldface.

| | 1 strong tape ; 0.25 N | | | | standard | | | | Total |
|---|---|---|---|---|---|---|---|---|---|
| | Cuticle broke | Detached | Not detached | Total | Cuticle broke | Detached | Not detached | Total | |
| ***D. ananassae*** | - | - | - | **-** | 0 | 29 | 1 | **30** | **30** |
| ***D. biarmipes*** | - | - | - | **-** | 0 | 48 | 1 | **49** | **49** |
| ***D. eugracilis*** | - | - | - | **-** | 2 | 29 | 0 | **31** | **31** |
| ***D. funebris*** | - | - | - | **-** | 1 | 36 | 0 | **37** | **37** |
| ***D. hydei*** | 5 | 36 | 14 | **55** | 3 | 4 | 23 | **30** | **85** |
| ***D. immigrans*** | 0 | 31 | 0 | **31** | 3 | 30 | 22 | **55** | **86** |
| ***D. kurseongensis*** | - | - | - | **-** | 0 | 18 | 0 | **18** | **18** |
| ***D. littoralis*** | 16 | 25 | 7 | **48** | 0 | 0 | 0 | **-** | **48** |
| ***D. malerkotliana*** | - | - | - | **-** | 2 | 32 | 0 | **34** | **34** |
| ***D. mauritiana*** | - | - | - | **-** | 0 | 23 | 0 | **23** | **23** |
| ***D. nannoptera*** | 8 | 13 | 5 | **26** | 8 | 19 | 17 | **44** | **70** |
| ***D. pachea*** | 7 | 11 | 7 | **25** | 1 | 8 | 22 | **31** | **56** |
| ***D. prostipennis*** | - | - | - | **-** | 0 | 28 | 0 | **28** | **28** |
| ***D. pseudoobscura*** | - | - | - | **-** | 0 | 30 | 0 | **30** | **30** |
| ***D. rhopaloa*** | - | - | - | **-** | 0 | 35 | 0 | **35** | **35** |
| ***D. simulans*** | - | - | - | **-** | 1 | 31 | 5 | **37** | **37** |
| ***D. suzukii*** | - | - | - | **-** | 1 | 55 | 0 | **56** | **56** |
| ***D. takahashii*** | - | - | - | **-** | 0 | 28 | 0 | **28** | **28** |
| ***D. tropicalis*** | - | - | - | **-** | 0 | 37 | 0 | **37** | **37** |
| ***D. virilis*** | 11 | 12 | 0 | **23** | 52 | 56 | 15 | **123** | **146** |
| ***D. yakuba*** | - | - | - | **-** | 2 | 41 | 3 | **46** | **46** |
| ***S. lebanonensis*** | - | - | - | **-** | 0 | 49 | 0 | **49** | **49** |
| ***Z. indianus*** | - | - | - | **-** | 1 | 28 | 0 | **29** | **29** |
| ***Z. lachaisei*** | 0 | 47 | 0 | **47** | 1 | 27 | 12 | **40** | **87** |
| **Total** | **47** | **175** | **33** | **255** | **78** | **721** | **121** | **920** | **1175** |

**Table S5. Correlation coefficients between adhesion and ecological parameters.** Rows indicate the developmental and ecological parameters, columns indicate the adhesion parameters. For each correlation, the corresponding r² and (uncorrected) p values from the linear regression are given, respectively in the first and second row; p values lower than 0.02 are in boldface.

| | Median detachment force | Median rigidity | Median deformation | Median pupal area | Median pupal shape | Median glue area | Median adhesion strength | Median glue investment |
|---|---|---|---|---|---|---|---|---|
| Number of cells per gland | 0.0082<br>0.79 | 0.037<br>0.57 | 0.0017<br>0.9 | 0.0073<br>0.87 | 0.028<br>0.75 | 0.0049<br>0.89 | 0.051<br>0.67 | 0.15<br>0.45 |
| Elevation | 9.3e-05<br>0.97 | 0.0083<br>0.69 | 0.015<br>0.6 | 0.00028 0.95 | 0.15<br>0.14 | 0.0026<br>0.86 | 0.00029<br>0.95 | 0.014<br>0.69 |
| Longitude | 0.12<br>0.13 | 0.11<br>0.14 | 0.2<br>0.04 | 0.015<br>0.66 | 0.036<br>0.48 | 0.19<br>0.12 | 0.022<br>0.62 | **0.41**<br>**0.013** |
| Latitude | 0.11<br>0.15 | 0.0018<br>0.86 | 0.11<br>0.15 | 0.035<br>0.49 | 0.14<br>0.15 | 0.051<br>0.44 | 0.026<br>0.58 | 0.078<br>0.34 |
| Mean temperature | 0.1<br>0.16 | 0.0094<br>0.68 | 0.13<br>0.11 | 0.072<br>0.32 | 0.24<br>0.055 | 0.04<br>0.49 | 0.00081<br>0.92 | 0.048<br>0.45 |
| Mean apparent temperature | 0.12<br>0.13 | 0.019<br>0.55 | 0.14<br>0.091 | 0.072<br>0.31 | 0.22<br>0.067 | 0.057<br>0.41 | 0.005<br>0.81 | 0.081<br>0.32 |
| Mean daylight duration | 0.14<br>0.094 | 0.0099<br>0.67 | 0.14<br>0.099 | 0.072<br>0.32 | 0.11<br>0.2 | 0.063<br>0.39 | 0.014<br>0.68 | 0.077<br>0.34 |
| Mean sunshine duration | 0.0019<br>0.85 | 0.12<br>0.12 | 4.7e-05<br>0.98 | 0.0095<br>0.72 | 0.084<br>0.28 | 0.024<br>0.6 | 0.013<br>0.69 | 0.081<br>0.32 |
| Mean precipitation | 0.14<br>0.092 | 0.17<br>0.066 | 0.18<br>0.057 | 0.056<br>0.38 | 0.015<br>0.65 | 0.21<br>0.099 | 0.0071<br>0.77 | **0.39**<br>**0.017** |
| Mean rain | 0.14<br>0.097 | 0.16<br>0.074 | 0.17<br>0.062 | 0.051<br>0.4 | 0.015<br>0.66 | 0.2<br>0.11 | 0.0077<br>0.77 | **0.38**<br>**0.019** |
| Mean snowfall | 0.00097<br>0.89 | 0.0051<br>0.76 | 7.8e-05<br>0.97 | 0.021<br>0.59 | 1.6e-05<br>0.99 | 0.0085<br>0.75 | 0.013<br>0.7 | 0.0028<br>0.86 |
| Mean wind speed | 0.16<br>0.077 | 0.037<br>0.4 | 0.13<br>0.11 | 0.095<br>0.24 | 0.037<br>0.48 | 0.19<br>0.12 | 0.011<br>0.72 | 0.26<br>0.063 |
| Mean wind gusts | 0.045<br>0.36 | 0.038<br>0.4 | 0.023<br>0.52 | 0.037<br>0.47 | 0.00095 0.91 | 0.05<br>0.44 | 0.17<br>0.15 | 0.07<br>0.36 |
| Mean shortwave radiation | 0.023<br>0.52 | 0.049<br>0.34 | 0.0095<br>0.67 | 0.037<br>0.48 | 0.051<br>0.4 | 0.0052<br>0.81 | 0.0014<br>0.9 | 0.037<br>0.51 |
| Mean reference evapotranspiration | 0.014<br>0.6 | 0.034<br>0.43 | 0.011<br>0.66 | 0.031<br>0.52 | 0.078<br>0.29 | 0.017<br>0.66 | 0.012<br>0.71 | 0.076<br>0.34 |
| Average humidity preference index | 0.18<br>0.057 | 0.0040.78 | 0.15<br>0.079 | 0.28<br>0.033 | 0.11<br>0.2 | 0.14<br>0.18 | 0.00066<br>0.93 | 0.046<br>0.46 |

### Supplementary files

**Available at DRYAD:**
**Final link (not active yet):**
**https://doi.org/10.5061/dryad.79cnp5j3j**

File S1. FileS1_dorsal_views_light_pupae.zip (1.99 GB). Dorsal views of light pupae. Compressed zip file of 1178 pupae pictures from the dorsal view (.tiff and .jpg). This file contains pictures of pupae manually selected for their light pupal case colour. The name of each image corresponds to the single pupa id, which is the date of adhesion test in format YYYYMMDD followed by a two-digit number which corresponds to the order in which pupae were processed on a given day. To find the corresponding species and associated information, please check Table “metadata_concatenate.csv” in the **'medata' folder of File S7,** whose column Sample_ID contains these single pupal identification numbers.

File S2. FileS2_dorsal_views_brown_pupae.zip (903.99 MB). Dorsal views of brown pupae. Compressed zip file of 389 pupae pictures from the dorsal view (.tiff and .jpg). This file contains pictures of pupae manually selected for their brown pupal case colour. The name of each image corresponds to the single pupa id (see legend of File S1).

File S3. FileS3_side_views.zip. Side views of all pupae (2.03 GB). Compressed zip file of 963 pupae pictures from the side view (.tiff and .jpg). The name of each image corresponds to the single pupa id (see legend of File S1) followed by "_side" or "_profile" or "-CantonS-profile" and the file extension.

File S4. FileS4_ventral_views.zip. Ventral views of all pupae (2.90 GB). Compressed zip file of 1403 pupae pictures from the ventral view (.tiff and .jpg). The name of most images corresponds to the single pupa id (see legend of File S1) followed by ‘_below or _down’ or "upsidedown' or "-CantonS-profile" and the file extension.

File S5. FileS5-ventral-views-measurements.zip (1.69 GB). Ventral views of all pupae with manual contours of the glue area (visible in ImageJ). Compressed zip file of 1276 pupae pictures (.tif). The name of each image is identical to the ones in File S4.

File S6. FileS6-ImageJ-Ilastic.zip (439.27 MB). Compressed zip file of a folder named "Pupa_contours" containing dorsal views with contours of the pupa area (visible in ImageJ) and ImageJ scripts (.ijm) and Ilastik training (.ilp) used to automatically measure pupal area and length on dorsal pictures. Scripts are named from 1 to 3 in the order of use. Script 1 is an ImageJ macro to convert the image files into a .tif format in order to be processed by Ilastik. Script 2 is a machine learning pixel classification made on Ilastik, respectively in the brown and light folders for dorsal pictures of brown and light pupae. Script 3 is an ImageJ macro to measure pupal area and length.

File S7. FileS7-csv-analyses.zip (543.33 MB). Compressed zip file of the input data, scripts (.R files) and output data (.pdf files).
